# A network of DZF proteins controls alternative splicing regulation and fidelity

**DOI:** 10.1101/2022.06.15.495552

**Authors:** Nazmul Haque, Alexander Will, Atlanta G. Cook, J. Robert Hogg

**Affiliations:** Biochemistry and Biophysics Center, National Heart, Lung, and Blood Institute, National Institutes of Health, Bethesda, MD, 20892, USA; Wellcome Centre for Cell Biology, School of Biological Sciences, University of Edinburgh, Max Born Crescent, Edinburgh, EH9 3BF, UK

**Keywords:** alternative splicing, DZF domain, ZFR, ILF2, ILF3, mutually exclusive splicing, RNA structure, zinc finger, double-stranded RNA binding domain, ultraconserved exon

## Abstract

Proteins containing DZF (<u>d</u>omain associated with <u>z</u>inc fingers) modules play important roles throughout gene expression, from transcription to translation. Derived from nucleotidyltransferases but lacking catalytic residues, DZF domains serve as heterodimerization surfaces between DZF protein pairs. Three DZF proteins are widely expressed in mammalian tissues, ILF2, ILF3, and ZFR, which form mutually exclusive ILF2-ILF3 and ILF2-ZFR heterodimers. Using eCLIP-Seq, we find that ZFR binds across broad intronic regions to regulate the alternative splicing of cassette and mutually exclusive exons. ZFR preferentially binds dsRNA *in vitro* and is enriched on introns containing conserved dsRNA elements in cells. Many splicing events are similarly altered upon depletion of any of the three DZF proteins; however, we also identify independent and opposing roles for ZFR and ILF3 in alternative splicing regulation. Along with widespread involvement in cassette exon splicing, the DZF proteins control the fidelity and regulation of over a dozen highly validated mutually exclusive splicing events. Our findings indicate that the DZF proteins form a complex regulatory network that leverages dsRNA binding by ILF3 and ZFR to modulate splicing regulation and fidelity.

## Introduction

Every step of eukaryotic gene expression involves collaboration between functional RNA elements and RNA-binding proteins (RBPs). Prototypical RBPs use one or more well-characterized RNA-binding domain (e.g. RNA recognition motifs, hnRNP K homology domains, and others) to contact specific RNA motifs. However, proteomic approaches have identified hundreds to thousands of novel RBPs, many of which lack recognizable RNA-binding protein folds (1, 2). Along with these new members of the RBP family, the regulatory and biochemical activities of many known RBPs remain unclear and require detailed investigation.

A vast number of RBPs participate in the regulation and execution of pre-mRNA splicing, a highly complex process that requires the coordinated assembly of hundreds of proteins on nascent pre-mRNAs (3). The catalytic steps of splicing are carried out by the spliceosome, a dynamic RNP enzyme that undergoes numerous conformational rearrangements throughout the splicing cycle (4–6). The core spliceosomal complexes are organized around small nuclear RNAs that directly recognize 5′ and 3′ splice sites and catalyze the splicing reaction. Each step in this process can be modulated by RBPs that are integrally or peripherally associated with the spliceosome. Recent structural studies of spliceosomal complexes have generated unprecedented insight into the organization of this RNP machine and illustrated the diversity of possible regulatory mechanisms available to exert splicing regulation (7–10). In parallel, genome wide studies of RBP interactions and functions have revealed that the complexity of the physical RNA-protein and protein-protein interaction networks is mirrored by the density of the regulatory networks comprising these proteins (11–13).

An intriguing example of an under-studied but evolutionarily conserved class of RBPs are a family containing DZF (<u>d</u>omain associated with <u>z</u>inc fingers) modules. DZF domains are derived from nucleotidyltransferases and retain a high degree of structural similarity to oligoadenylate and cyclic GMP-AMP synthetases; however, they lack key catalytic residues and instead serve as protein-protein interaction domains (14). Humans express only five DZF-encoding genes, of which the most extensively studied are ILF2 (also known as NF45) and ILF3 (also known as NF90/NF110), which heterodimerize via their DZF domains (14, 15). The ILF2 and ILF3 genes have been implicated in almost every aspect of RNA biology, including alternative splicing, miRNA processing, circular RNA production, translation, RNA adenosine-to-inosine editing, and transcription (16–26). The third widely expressed DZF protein is zinc finger RNA-binding protein (ZFR), shown to be an important regulator of both alternative splicing and RNA editing in humans and *Drosophila* (23, 27–29). ILF2, ILF3, and ZFR all exhibit broad expression throughout human and mouse tissues (Figure S1), and ILF2 and ILF3 have been estimated by integrative proteomic analysis to be one order of magnitude more abundant than ZFR (30). The remaining human DZF proteins, the ILF3 paralog SPNR and the ZFR paralog ZFR2, are restricted to specific cell types (31).

Biochemical and structural studies of ILF2-ILF3 heterodimerization show that the DZF domains of ILF3 and ZFR engage in mutually exclusive interactions with the ILF2 DZF domain (Figure 1A and B; 14). ILF2 is required for stable accumulation of ILF3 protein and may modify its RNA binding activities (32, 33). Even though ZFR-ILF2 is the evolutionarily older complex, found in flies and worms (Figure S2A; 34), the effects of ILF2 on ZFR function have not been reported. Interestingly, ILF3 and ZFR contain distinct RNA-binding modules: ZFR uses three highly conserved C2H2 zinc fingers to bind RNA with unknown specificity, while ILF3 preferentially binds long dsRNAs via two dsRBDs (Figure 1A; 34). This arrangement of physical interactions raises the possibility that the human DZF proteins form a functional regulatory network modulating multiple aspects of RNA biology, including RNA editing and RNA splicing.

**Figure 1.**
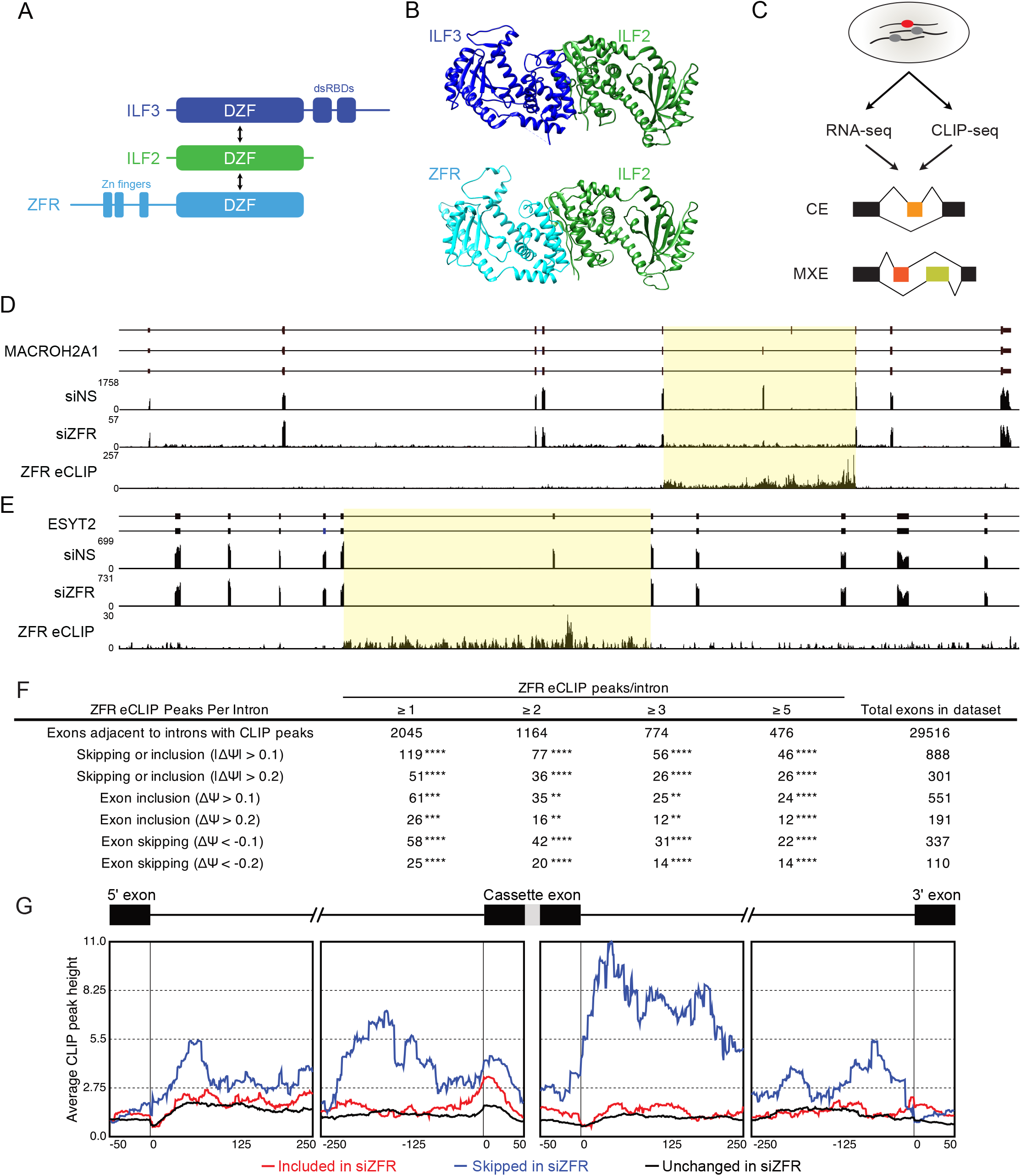
ZFR binds large intronic domains flanking regulated exons. A. Schematic of ZFR, ILF2, and ILF3 protein features and heterodimerization through DZF domains. B. Top, crystal structure of the heterodimer interface between ILF2 and ILF3 DZF domains (PDB 4AT7; 14). Bottom, Alphafold-Multimer prediction of the structure of the heterodimer formed by the ZFR and ILF2 DZF domains (69). C. Scheme for using CLIP-seq and RNA-seq to investigate interactions between ZFR and pre-mRNAs undergoing CE or MXE alternative splicing. D. ZFR eCLIP and ZFR and control knockdown RNA-seq read histograms from MACROH2A1 are shown below gene models corresponding to MXE splicing products. Direction of transcription is left to right. Introns flanking ZFR-regulated exons are indicated by yellow shading. E. ZFR eCLIP and ZFR and control knockdown RNA-seq read histograms from ESYT2 are shown below gene models corresponding to CE splicing products. Direction of transcription is left to right. Introns flanking ZFR-regulated exons are indicated by yellow shading. F. Table comparing the incidence of ZFR eCLIP peaks in introns adjacent to exons identified as significantly regulated in rMATS analysis of siZFR vs. siNT RNA-seq. Empirical *P* values were determined by permutation testing with the RegioneR package (** *P* < 0.01; *** *P* < 0.001; **** *P* < 0.0001; 70). G. Metagene analysis of ZFR eCLIP peak position and frequency relative to cassette exons skipped or included upon siZFR treatment, generated using rMAPS software (71).

Here, we investigate the physical and functional interactions of ILF2, ILF3, and ZFR. Beginning with the first transcriptome-wide CLIP-seq study of ZFR-RNA interactions, we find that ZFR binds large domains in introns flanking the cassette and mutually exclusive exons (CEs and MXEs, respectively) it regulates. We find that ZFR binds dsRNA via its zinc fingers *in vitro* and binds and regulates pre-mRNAs containing conserved secondary structural elements in cells. RNA-seq studies of cells depleted of ZFR, ILF2, or ILF3 reveal a complex functional relationship among the proteins. While many splicing events are similarly affected by depletion of each of the three DZF proteins, we identify CEs and MXEs that are antagonistically regulated by ILF3 and ZFR. Highlighting the importance of DZF proteins in MXE splicing, we identify several instances in which MXE splicing fidelity is impaired upon DZF protein depletion, resulting in aberrant skipping or fusion of MXEs.

## Results

### ZFR interacts with long stretches of intronic sequences

We have previously shown that ZFR depletion causes dysregulated splicing of many CEs and MXEs, but how ZFR recognizes its pre-mRNA targets is unknown (28). We therefore performed UV crosslinking and immunoprecipitation followed by high-throughput sequencing (CLIP-Seq) to determine the relationship between ZFR binding and splicing regulatory function throughout the transcriptome (Figure 1C). For these experiments, we integrated FLAG-tagged ZFR cDNA into HEK-293 Flp-In T-REx cells, allowing doxycycline-inducible FLAG-ZFR expression at levels closely matching those of endogenous ZFR (Figure S2B). FLAG-tagged GFP-expressing lines were used as a control to confirm the absence of nonspecific RNA recovery. To identify ZFR binding sites, we used the enhanced CLIP (eCLIP) protocol, modified for sensitive detection of RNA-protein complexes in SDS-PAGE by inclusion of an infrared dye on the 3′ adapter oligonucleotide ligated to RBP-bound RNA fragments (35, 36). ZFR eCLIP libraries were highly enriched for intronic sequences, consistent with the established role of ZFR in alternative splicing (Figure S2C). Exonic ZFR binding was similarly distributed across coding and 3′UTR sequences, while we observed only low levels of binding to 5′UTRs (Figure S2D).

Inspection of previously characterized ZFR targets revealed that ZFR frequently binds throughout large intronic stretches flanking ZFR-regulated exons (Figure 1D and E; 28). This pattern was observed both for ZFR-regulated MXEs (e.g. MACROH2A1, Figure 1D) and CEs (e.g. ESYT2, Figure 1E; PAM, Figure S2E). We previously reported two modes of regulation of MXEs by ZFR: skipping of both potential MXEs or switching between MXEs (28). Using eCLIP, we observed direct binding of ZFR to pre-mRNAs undergoing both types of ZFR-dependent MXE regulation. Direct regulation of MXE skipping was exemplified by ZFR binding to a 14 kb segment of MACROH2A1 pre-mRNA containing mutually exclusive exons that are both efficiently skipped in the absence of ZFR (Figure 1D). Similarly, we observed broad regions of ZFR eCLIP enrichment near receptor tyrosine kinase FYN MXEs, the usage of which dramatically switches upon ZFR depletion (Figure S2F).

To investigate the transcriptome-wide relationship between ZFR binding and splicing regulation, we examined the occurrence of ZFR eCLIP peaks in introns flanking CEs affected by ZFR depletion from HEK-293 cells (Figure 1F; Table S1). In this analysis, we stratified splicing events according to the effect of ZFR depletion (CE skipping vs. inclusion), the magnitude of the ZFR-induced change in inclusion, and the number of ZFR eCLIP peaks identified in the introns flanking the regulated exon. We observed a significant overlap between sites of ZFR binding and function in all cases, supporting a direct role for ZFR in promoting both exon skipping and inclusion. Metagene analyses also implicated broad regions of ZFR binding in splicing regulation, with the highest levels of ZFR CLIP signal observed surrounding exons skipped in the absence of ZFR (Figure 1G).

### ZFR regulates detained intron splicing

While most splicing regulators are thought to function by binding specific sites in or near exons, ZFR binding throughout introns is reminiscent of the observation that SRSF3 and SRSF7 bind throughout certain inefficiently spliced introns (37). We further investigated the links among ZFR binding, function, and intron retention, focusing on introns previously identified as detained introns (DIs), meaning that they persist in nuclear mRNAs following the completion of other splicing and processing events (38). Messages containing DIs can attain multiple fates, including post-transcriptional splicing, nuclear degradation, or export to the cytoplasm. Transcriptome-wide, we found a high degree of overlap between DI and ZFR CLIP peaks (Figure 2A; Figure S3A). The physical association of ZFR with DI was accompanied by a strong functional relationship, as ZFR exhibited a propensity to regulate splicing of exons flanked by DIs (Figure 2B; Figure S3B-F).

**Figure 2.**
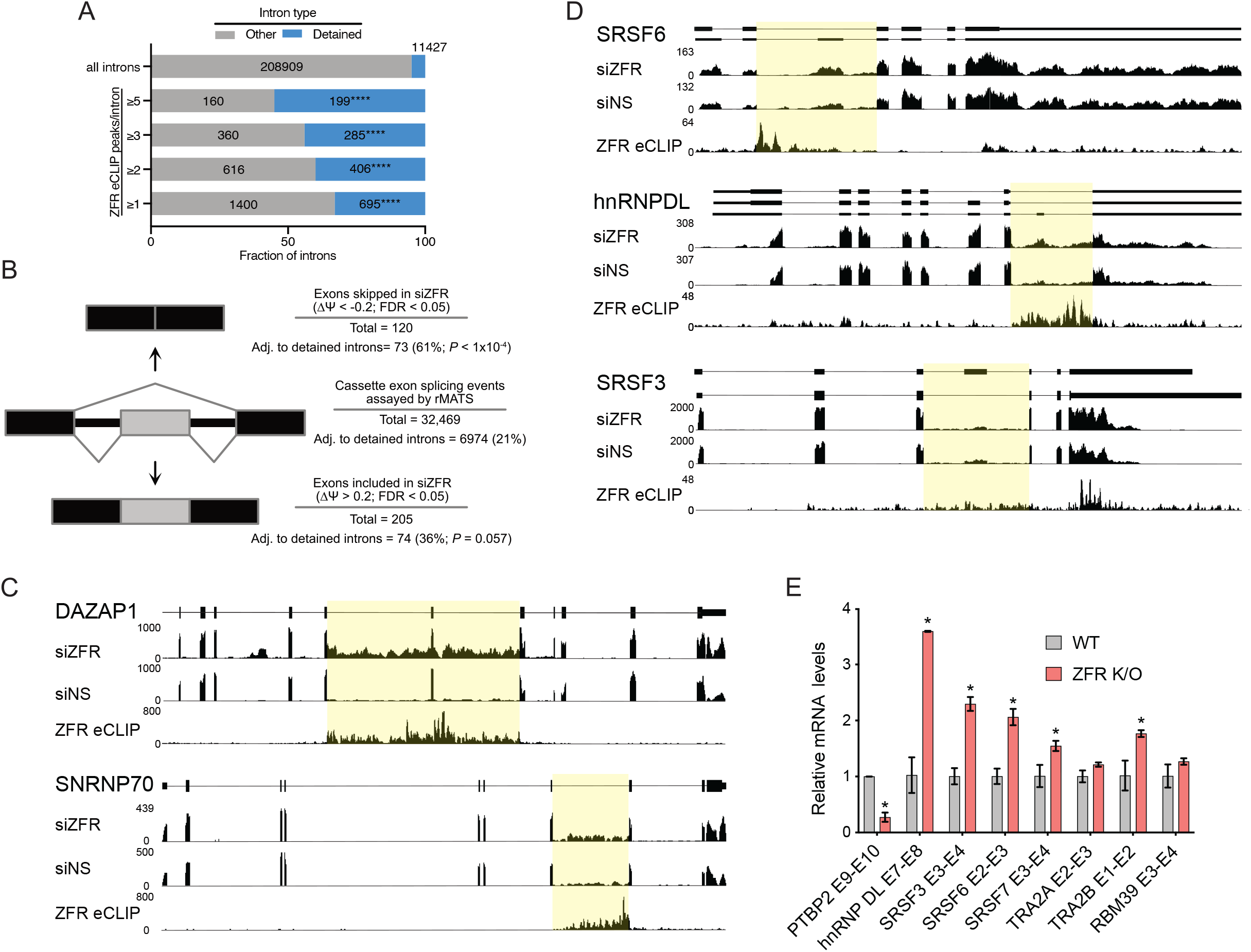
ZFR binds detained introns and regulates the splicing of adjacent exons. A. Table of the incidence of ZFR eCLIP peaks in detained introns identified in (38). Empirical *P* values were determined by permutation testing with the RegioneR package (**** *P* < 0.0001; 70). B. Schematic illustrating the relationship between detained introns and ZFR-mediated skipping (top) and inclusion (bottom) of CEs. Empirical *P* values were determined by permutation testing with the RegioneR package (70). C. ZFR eCLIP and ZFR and control knockdown RNA-seq read histograms from the indicated genes that undergo increased intron retention in the absence of ZFR. D. ZFR eCLIP and ZFR and control knockdown RNA-seq read histograms from the indicated genes containing ultraconserved exons. Gene models corresponding to alternative splice isoforms are shown. E. Isoform specific RT-qPCR of RNA isolated from ZFR knockout or parental control HEK-293 cells to assess alternative splicing of ultraconserved exons. Relative levels of mRNA isoforms containing the indicated exon junctions were determined by normalization to GAPDH mRNA. Statistical significance was determined by two-way ANOVA, with Sidak’s correction for multiple comparisons (n = 3; error bars indicate ±SD; *P* < 0.05).

Although our data suggest a role for ZFR in regulating splicing of exons adjacent to detained introns, we did not observe a widespread impact of ZFR depletion on the steady-state levels of DIs themselves (Table S1). We did find rare cases in which reads derived from DIs were highly elevated in the absence of ZFR, including introns in DAZAP1 and SNRNP70 (Figure 2C; 38). In both cases, we observed strong ZFR eCLIP signals, suggesting a direct role for ZFR in promoting removal or turnover of these introns. Despite these examples, we conclude that ZFR frequently regulates the outcomes of post-transcriptional splicing events but does not systematically affect the steady-state accumulation of inefficiently spliced introns.

### ZFR regulates splicing of ultraconserved exons

The ZFR locus and several ZFR target genes contain exons embedded in ultraconserved elements (UCEs), defined as sequence blocks of 200 bp or greater that are 100% identical among the human, mouse, and rat genomes (39). Ultraconserved exons are associated with alternative splicing and nonsense-mediated decay (AS-NMD) of transcripts encoding important splicing regulators (40, 41). Recent analyses have identified a highly complex network of interactions among UCE-containing splicing factors, whereby proteins produced from UCE-containing genes frequently go on to regulate splicing of their own and other UCE exons (42). The status of ZFR as a UCE-containing gene and several examples of strong ZFR eCLIP signal on or near UCEs (e.g. SRSF3, SRSF6, and hnRNP DL; Figure 2D) thus prompted us to further investigate its role in modulating UCE-associated splicing. We found that abolishing full-length ZFR expression by CRISPR/Cas9-mediated disruption of exons 8 and 9 in HEK-293 Tet-off cells caused altered splicing of several UCE exons, including those from the PTBP2, HNRNPDL, SRSF3, SRSF6, SRSF7, TRA2A, TRA2B, and RBM39 transcripts (Figure 2E; Figure S3G). In each case, ZFR depletion led to enhanced production of the NMD-sensitive isoform, suggesting that ZFR maintains cellular levels of splicing regulators by promoting UCE-associated splicing events that produce stable mRNAs encoding full-length proteins.

### ZFR preferentially binds dsRNA *in vitro*

Analysis of ZFR eCLIP data indicated specific binding to intronic regions flanking regulated exons but did not generate immediate insight into RNA sequence or structural determinants of ZFR binding. To directly analyze the RNA binding properties of ZFR, we produced recombinant ZFR containing the zinc finger motifs (ZFR_long_; residues 317-1074) or a C-terminal fragment of ZFR containing the DZF domain (ZFR_DZF_; residues 725-1074) in *E. coli.* To promote proper folding and solubility, we co-expressed the ZFR variants with a near full-length ILF2 protein (residues 29-390), forming a stable ZFR-ILF2 heterodimer via the two proteins’ DZF domains (Figure S4A and B). We first asked whether the zinc fingers of ZFR are required for RNA binding by performing electrophoretic mobility shift assays (EMSAs) with ZFR_long_/ILF2 and ZFR_DZF_/ILF2 and a either single-stranded U_20_ ssRNA (Figure 3A) or a 20 bp U:A dsRNA (Figure 3B). The ZFR_long_/ILF2 heterodimer exhibited weak binding to the poly-U ssRNA, while the heterodimer lacking the ZFR zinc fingers showed no capacity to interact with this substrate (Figure 3A). ZFR_long_/ILF2 bound substantially more tightly to the U:A dsRNA, again in a manner dependent on the ZFR zinc fingers (Figure 3B).

**Figure 3.**
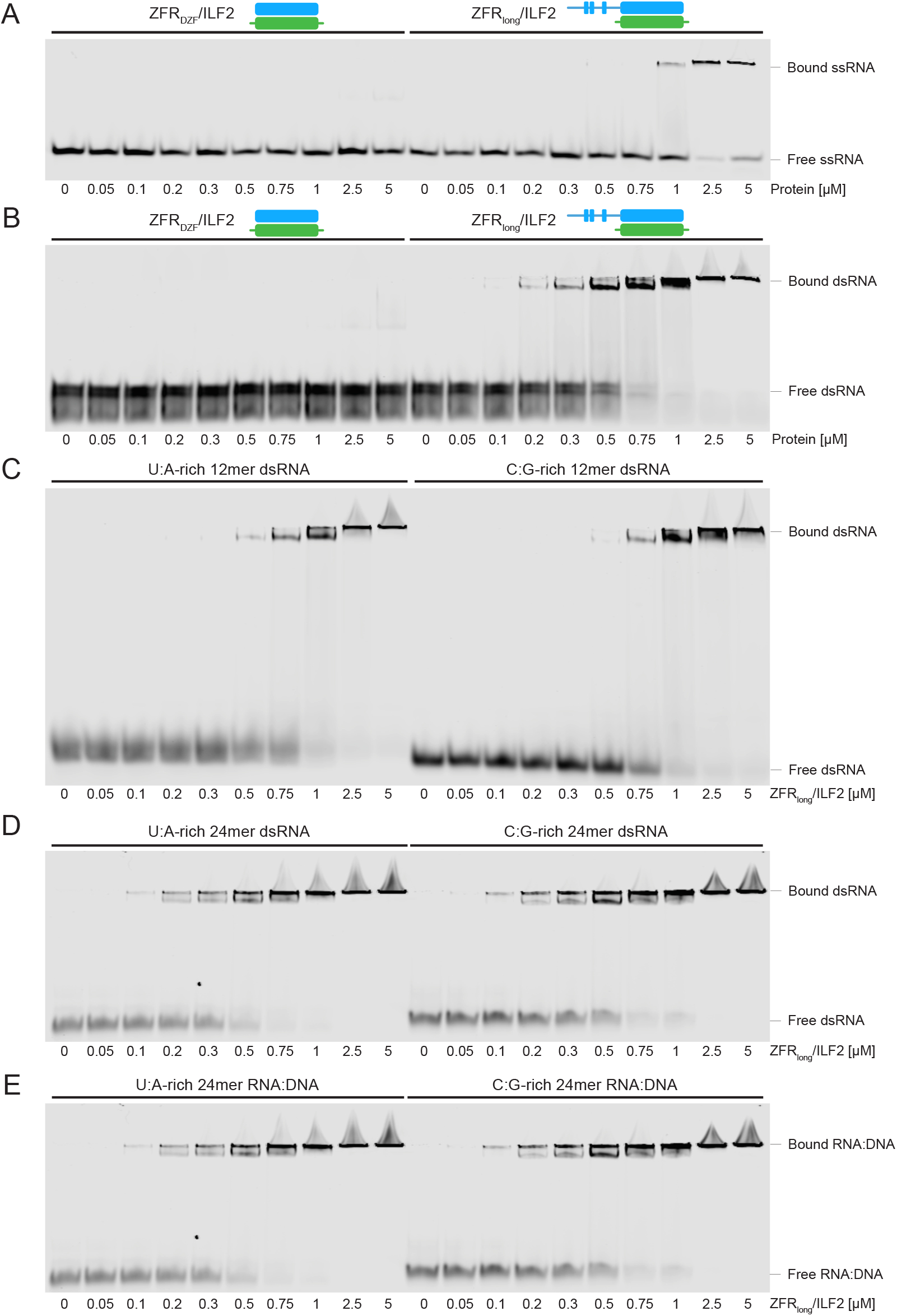
ZFR prefers dsRNA or DNA-RNA hybrids over ssRNA. A. ZFR variants containing (ZFR_long_) or lacking zinc-fingers (ZFR_DZF_) were co-expressed and purified with ILF2. Binding of the indicated ZFR/ILF2 protein concentrations to fluorescently labeled single-stranded 12-mer oligo-U RNA was assayed by EMSA. B. As in A, with dsRNA probe formed from 12-mer oligo-U and oligo-A sequences. C. As in A, with U-rich (left) and C-rich dsRNA 12-mer probes. D. As in C, with 24-mer probes. E. As in D, with U-rich and C-rich 24-mer RNAs duplexed to complementary A-rich and G-rich DNAs.

To further investigate ZFR binding to dsRNA, we assayed ZFR_long_/ILF2 binding to 12 bp AU-rich or GC-rich dsRNAs. ZFR did not exhibit a preference between the distinct 12 bp duplex sequences (Figure 3C, K_d_ ∼ 0.8 μM). ZFR bound more tightly to 24 bp dsRNAs containing AU- or GC-rich sequences (Figure 3D, K_d_ ∼ 0.3 μM) than to the 12-mer substrates but again did not show differential binding on the basis of sequence composition. Indicative of a general propensity to bind A-form duplex nucleic acids, the affinity of ZFR_long_/ILF2 for RNA:DNA hybrids was similar to that observed for dsRNAs (compare Figure 3E with Figure 3D). Together, our data suggest that ZFR binds to dsRNA or DNA:RNA hybrids with high affinity but little apparent sequence specificity. The presence of multiple low-mobility complexes in the EMSA analyses, particularly on the 24mer substrates, suggests either multiple possible conformations or oligomerization of ZFR/ILF2 on A-form duplex nucleic acids.

We also employed a sequencing-based approach, RNA Bind-n-Seq (RBNS), to characterize the RNA-binding properties of ZFR (43, 44). To perform RBNS, we incubated a library of *in vitro* transcribed RNAs containing 40 nt long random sequences flanked by primer binding sites for sequencing library preparation with purified ZFR_long_/ILF2, using ZFR_DZF_/ILF2 and recombinant polypyrimidine tract binding protein 1 (PTBP1) as controls. The recombinant proteins were affinity purified on magnetic nickel beads, and sequences enriched or depleted from the starting pool were monitored by high-throughput sequencing. Unexpectedly, the most highly enriched sequences in ZFR_long_/ILF2 RBNS were complementary to a sequence in the 3′ PCR primer-binding site flanking the 40 nt random sequence (Figure S5A and B; Table S2). Because the primer-binding sequence was present in all molecules in the library, complementary sequences within the randomized region would be the most abundant dsRNA-derived sequences in the pool (Figure S5C). The ZFR_long_/ILF2 results contrasted with the results of the PTBP1 RBNS control, in which CU-rich sequences in ssRNA context were preferentially recovered, matching previous findings using RBNS and other approaches (Figure S5D; 43). Together, these results suggest that the preference of ZFR for double-stranded RNA caused it to recover sequences with base-pairing potential with the PCR primer binding sites (Figure S5C). Because these results are biased by the sequence identity of the primer binding sites, we cannot use them to make conclusions regarding ZFR sequence specificity.

### DZF proteins form an intricate regulatory network

ILF2 uses its DZF domain to heterodimerize with either ZFR or the dsRBD-containing ILF3 protein, raising the question of why vertebrates have evolved two ILF2-interacting proteins that bind dsRNA via structurally unrelated protein modules. To better understand the physical and functional relationships among the three DZF domain proteins, we first generated cell lines stably expressing FLAG-tagged ILF2 and ILF3 proteins, with FLAG-GFP as a control. Consistent with previous findings (14), immunoaffinity purified FLAG-ILF2 complexes contained endogenous ILF3 and ZFR, while FLAG-ILF3 only co-purified with endogenous ILF2, not ZFR (Figure 4A).

**Figure 4.**
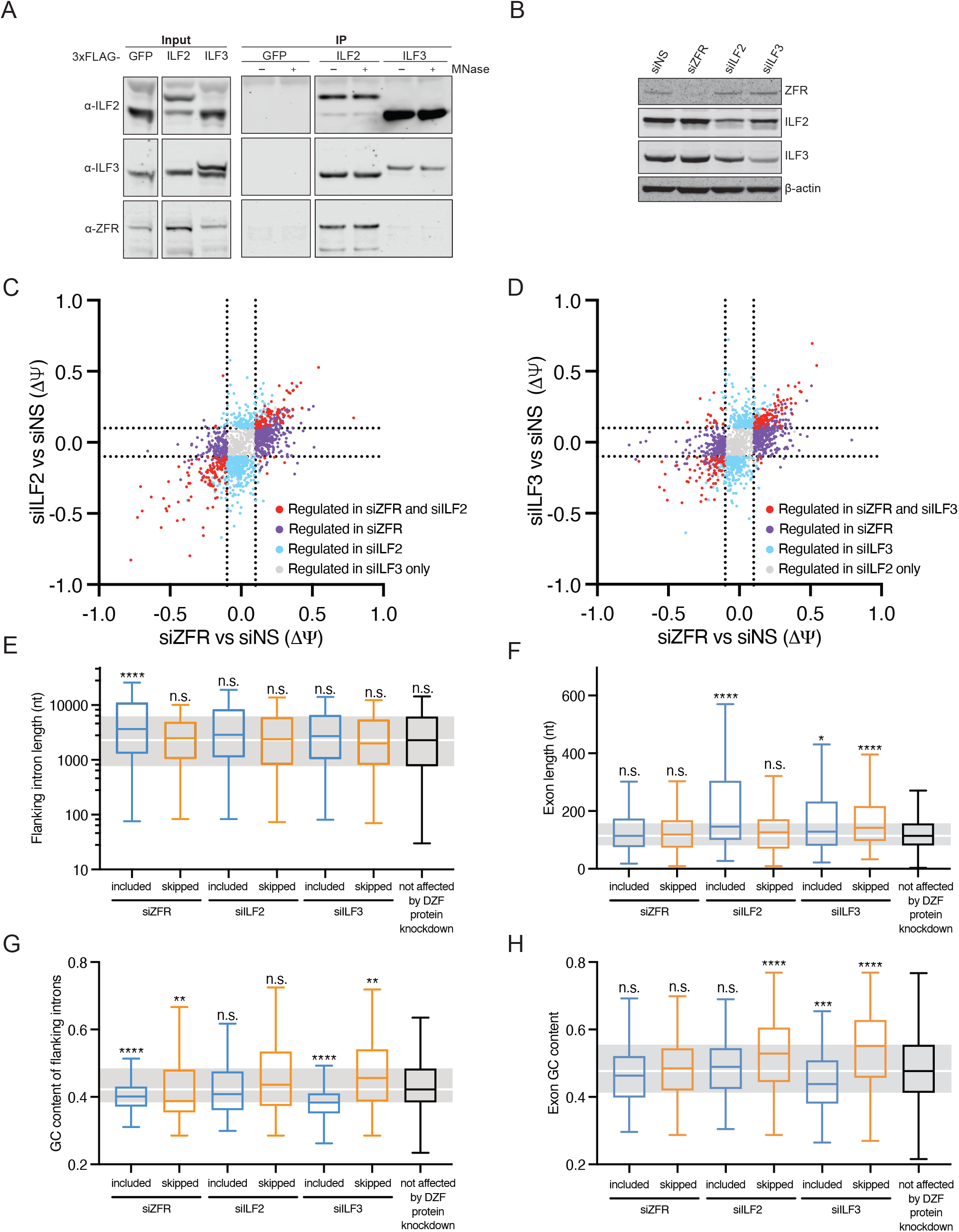
DZF proteins ILF2, ILF3, and ZFR form a complex regulatory network. A. Immunoblot of input (left) and bound (right) fractions from immunoaffinity purification of the indicated stably expressed FLAG-tagged proteins in the presence (+) and absence (-) of MNase. Membranes were probed with antibodies recognizing the endogenous ILF2, ILF3, and ZFR proteins as indicated. All panels are derived from identical exposures of the same membrane. B. Immunoblot of whole-cell extracts from cells treated with the indicated siRNAs, using antibodies against the endogenous ZFR, ILF2, ILF3, and β-actin. C. Scatterplot comparing response of CE events meeting rMATS read count cutoffs and regulated in at least one DZF protein knockdown condition (|ΔѰ| > 0.1 and FDR < 0.05) to depletion of ZFR or ILF2. Dotted lines indicate ΔѰ ± 0.1. D. As in C, comparing the effects of ZFR and ILF3 depletion. E. Box plot indicating the distribution of lengths of introns flanking of exons regulated by depletion of the indicated DZF proteins (|ΔѰ| > 0.1 and FDR < 0.05 in rMATS analysis) or exons unaffected by any DZF protein knockdown. Boxes indicate interquartile ranges, center lines indicate medians, and whiskers indicate Tukey ranges. Gray regions correspond to interquartile ranges in the negative control distribution of introns adjacent to unaffected exons. Statistical significance was determined by two-tailed Kruskal-Wallis test, with Dunn’s correction for multiple comparisons. F. As in D, with lengths of exons in the indicated classes. G. As in D, with GC content of introns adjacent to exons in the indicated classes. H. As in D, with GC content of exons in the indicated classes.

We next used specific siRNAs to individually deplete ZFR, ILF2, or ILF3 from HEK-293 Tet-off cells. Analysis of ZFR, ILF2 and ILF3 mRNAs and proteins in the knockdown cells revealed further evidence for a dense regulatory network among the three DZF proteins. In qRT-PCR analyses of knockdown cells, ZFR mRNA levels were efficiently reduced by siZFR treatment but were induced approximately 2.5-fold by ILF2 knockdown (Figure S6A). ILF3 knockdown did not affect endogenous ZFR mRNA levels, and ILF2 mRNA levels were only substantially affected by the ILF2 siRNA (Figure S6B). Similar to ZFR, siILF2 treatment caused enhanced ILF3 mRNA expression (Figure S6C). Along with ILF2-mediated regulation of ILF3 and ZFR mRNA expression, we observed that levels of both ILF2 and ILF3 proteins were reduced upon treatment with either siILF2 or siILF3, as previously reported (Figure 4B). ZFR protein levels were not affected by knockdown of either ILF2 or ILF3, and ZFR knockdowns did not decrease ILF2 or ILF3 levels. However, a version of ZFR in which the DZF domain residues predicted to be required for ILF2 binding were mutated was poorly expressed (E1008R; R1019E), suggesting that ZFR may also rely on ILF2 for stability in cells, but that residual ILF2 remaining after siILF2 treatment is sufficient to maintain ZFR protein levels (Figure S6D).

### Regulation of cassette exons by ZFR, ILF2, and ILF3

We next investigated the functional relationships among ZFR, ILF2, and ILF3 by analyzing transcriptome-wide splicing alterations in cells depleted of each protein. Consistent with the physical interdependencies discussed above, depletion of each DZF-domain protein had broadly similar impacts on global regulation of CE splicing patterns (Figure 4C and D; Figure S6E; Table S1). Of 734 CE splicing events affected by siZFR (defined as |ΔѰ| > 0.1 with rMATS FDR < 0.05), 326 were regulated by depletion of ILF2 and/or ILF3. Of these, 94 were regulated in response to depletion of all three DZF proteins (Figure S6E).

An important caveat to these studies is that network of DZF protein interactions complicates interpretation of the splicing changes in ZFR, ILF2, and ILF3 knockdown conditions. While our ZFR CLIP studies suggest a strong relationship between ZFR binding and splicing regulation, some observed changes in splicing could be due either to direct effects on the target DZF protein or to indirect effects on the other DZF proteins. Such indirect effects could be complex: for example, ILF3 depletion could cause the inverse phenotype to ZFR depletion either because the proteins directly antagonize each other or because ILF3 depletion relieves competition between ILF3 and ZFR for ILF2 binding. Conversely, reduced ILF2 protein stability upon ILF3 depletion could reduce the pool of ILF2 available for heterodimerization, indirectly causing a reduction in ZFR activity.

Despite the physical and regulatory interdependence of the DZF proteins and the high degree of overlap among CEs regulated by ZFR, ILF2, and ILF3, we uncovered interesting differences among the splicing events perturbed by depletion of the three proteins (Figure 4E-H). Exons included upon siZFR treatment were associated with long 5′ and 3′ flanking introns (median 5′ and 3′ intron length of 4513 nt and 3259 nt, respectively, for exons included in siZFR, *versus* 2400 nt and 2203 nt for exons not regulated by any of the DZF proteins), while the distribution of intron sizes among ILF2 and ILF3 targets was not distinguishable from that of the overall set of introns represented in the dataset (Figure 4E). Conversely, exons included in siILF2 (mean length 146 nt) and both included (128 nt) and excluded (142 nt) in siILF3 were significantly longer than those in the overall set of exons assayed (114 nt; Figure 4F).

Along with differences in the lengths of exons and introns, we also observed distinct nucleotide compositions for ZFR and ILF3 targets. In mammals, long introns tend to have low GC content and a high differential in exon-intron GC content, while short introns have higher GC content and a low exon-intron GC content differential (45). Consistent with ZFR-mediated regulation of exons associated with long introns, exons included (median 46.3% GC vs. 47.7% GC in exons not affected by DZF protein depletion) and skipped (median 48.5%) as a result of ZFR knockdown were embedded in introns with low GC content (median 38.8% GC for included; median 40.1% GC for skipped) relative to that of the set of control introns (42.2% GC; Figure 4G and H). Interestingly, ILF3 knockdown caused inclusion of low-GC exons (median 44% GC vs. 47.7% GC in control exons) in low-GC content introns (median 38.3% GC) and skipping of high-GC exons (55.1% GC) in high-GC introns (median 45.6% GC; Figure 4G and H).

In addition, we identified several splicing events that were significantly altered in opposite directions by ZFR and ILF3 knockdown (Figure 4D). Subsequent qPCR analyses validated several cases of opposing ILF3 and ZFR regulation of cassette exons, including exons from LRIG2 (Figure S7A), PPP4R1 (Figure S7B), and SLC38A1 (Figure S7C). On each of these targets, ILF2 knockdown phenocopied ILF3 knockdown, despite the presumed requirement of ILF2 for ZFR function.

### ZFR binds to conserved secondary structures across the transcriptome

A recent report identified conserved structured regions throughout the transcriptome, designated pairs of conserved complementary regions (PCCRs; 46). These regions, determined by computational methods and corroborated by experimental data, are associated with alternative splicing and RNA editing. Because ZFR and ILF3 bind dsRNA and, along with ILF2, regulate both splicing and editing, we investigated a potential link between PCCRs and DZF protein binding and function.

We first asked whether PCCRs were enriched in introns flanking exons affected by DZF protein depletion. With the set of introns flanking exons evaluated in our rMATS analysis as a search space, we evaluated co-occurrence of PCCRs and DZF protein alternative splicing targets across the transcriptome. We found that PCCRs were enriched in introns flanking exons skipped or included upon ZFR knockdown, with over half of the most strongly ZFR-regulated exons (|ΔΨ| > 0.2) flanked by one or more PCCRs, *versus* 31% of all exons not affected by ZFR depletion (Figure 5A). Interestingly, ZFR targets were enriched for multiple flanking PCCRs, with 26% of included exons and 24% of skipped exons residing adjacent to introns with six or more PCCRs, relative to just 12% of exons in the total population. We also observed enrichment of PCCRs in pre-mRNAs strongly affected by ILF2 depletion (|ΔΨ| > 0.2), but the corresponding set of ILF3 targets were not significantly associated with PCCRs (Figure 5A).

**Figure 5.**
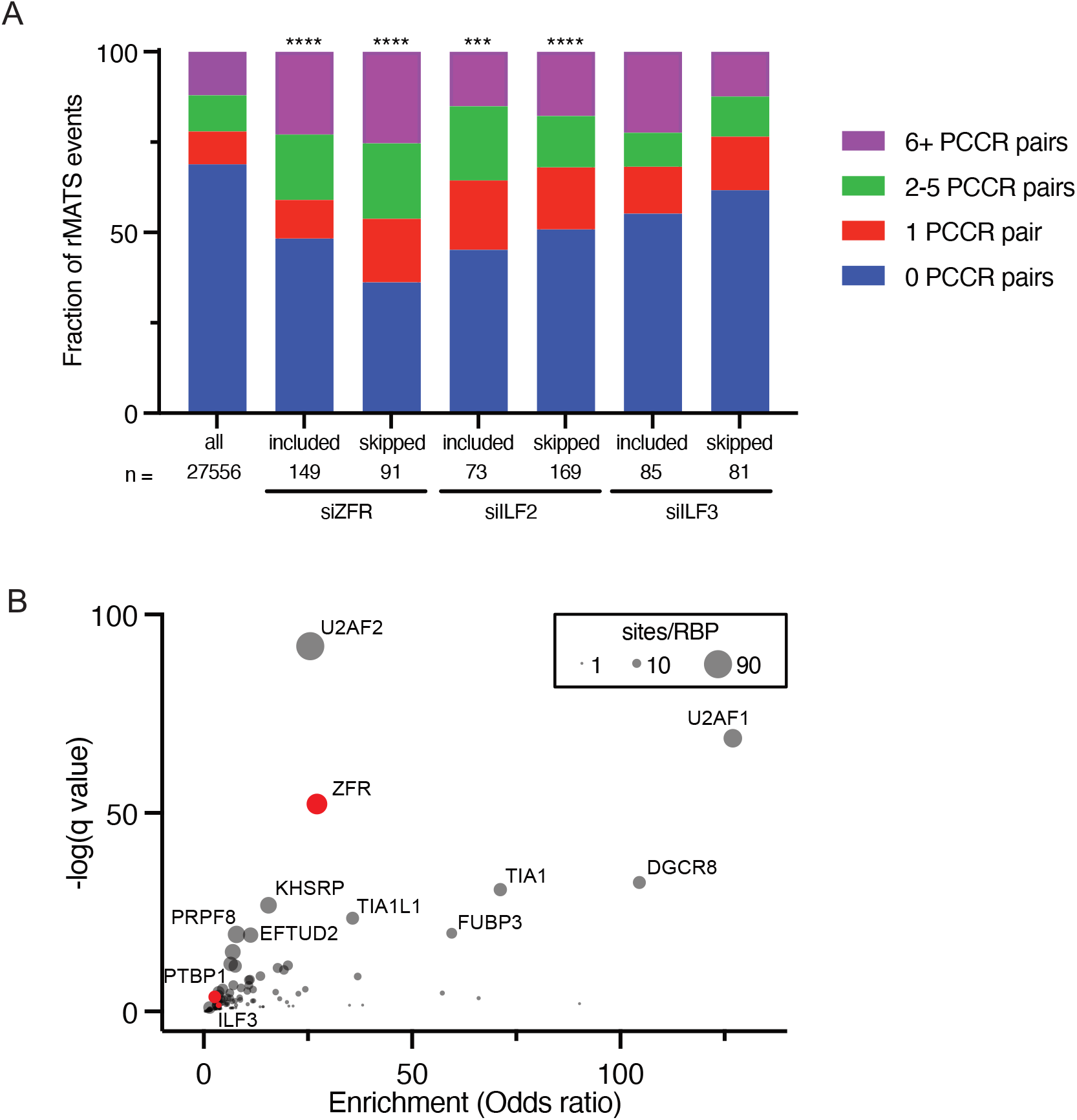
ZFR binds to and functions on introns containing conserved dsRNA elements. A. Correspondence between CEs skipped or included in siZFR, siILF2, or siILF3 treatment relative to siNS controls (FDR < 0.05 and |ΔѰ > 0.2| in rMATS analysis) and exons adjacent to introns containing the indicated numbers of conserved sequences predicted to form dsRNA (PCCR pairs; 46). *P* values indicate significant enrichment of introns with one or more PCCRs among the indicated response categories and were determined by Chi-square test (\**P* < 0.05; \*\**P* < 0.01; \*\*\**P* < 0.005; **** *P* < 0.0005). B. Correspondence between eCLIP peaks identified in ZFR and PTBP1 eCLIP (red circles; this study) and ENCODE eCLIP studies (black circles except for ILF3 in red; 48) and predicted conserved intronic dsRNA elements (46). Circle sizes correspond to the number of PCCR sites occupied by the corresponding RBPs. Odds ratios and q values were computed using the LOLA package (47).

We also investigated whether ZFR and ILF3 eCLIP peaks were enriched on PCCRs. For this analysis, we compared our ZFR eCLIP data with PTBP1 eCLIP data prepared in parallel, along with eCLIP data from the ENCODE consortium (47, 48). Among all RBPs assessed, ZFR was one of the most highly and significantly enriched on PCCRs, while in-house and ENCODE PTBP1 and ENCODE ILF3 eCLIP showed no enrichment for PCCRs (Figure 5B). Together, these data indicate that ZFR may function in concert with conserved double-stranded regions to select targets for pre-mRNA splicing regulation.

### Effects of DZF proteins on MXE splicing regulation

We previously observed that ZFR depletion preferentially affects splicing of mutually exclusive exons (28). rMATS identified 148 putative MXE alternative splicing events from 129 genes regulated by one or more DZF protein (|ΔѰ| > 0.1 with rMATS FDR < 0.05; Table S1). As observed for CEs, there was a high degree of overlap among putative MXEs affected by ZFR, ILF2, and ILF3 knockdown, with the strongest effects elicited by ZFR and ILF2 depletion (Figure S8A and B). However, the complex architecture of MXEs means that identification approaches based on short reads are subject to high rates of both false-positives and false-negatives (49). Manual inspection of rMATS results corroborated the identification of many genuine instances of DZF-mediated MXE regulation but also revealed extensive false-positive MXE calls by rMATS. To assemble a catalog of genuine MXEs regulated by the DZF proteins, we therefore validated rMATS results with existing MXE databases and manual inspection.

Several attempts to catalog human MXEs have been made (49–52), from which we assembled a master list of 810 genes with previously annotated MXEs (Table S3). Of the 810 genes in this list, 229 met our stringent rMATS junction read count cutoffs, 22 of which were identified as significant MXE targets of one or more DZF protein by rMATS (13 regulated in siZFR, 16 regulated in siILF2, and 9 regulated in siILF3; note that some MXE events were regulated by multiple DZF proteins). Supporting the validity of this approach, almost all of the significantly regulated events were represented in at least two datasets. Among 58 genes that were represented in two or more MXE datasets and met our rMATS junction read cutoffs, 16 (28%) were significantly regulated by at least one DZF protein (12 regulated in siZFR, 13 regulated in siILF2, and 6 regulated in siILF3; note that some MXE events were regulated by multiple DZF proteins).

To verify DZF-regulated MXEs, we used a manual inspection approach to distinguish between genuine MXE splicing and CE skipping/inclusion events. Our manual inspection criteria required junction-spanning reads connecting the 5′ and 3′ splice sites of two or more putative mutually exclusive exons to flanking constitutive exons (Figure 6A and B; e.g. DNM2, Figure 6C). We included genes in which there was substantial skipping of both putative MXEs and genes in which both putative MXEs were observed in a subset of transcripts. This approach eliminated many splicing events that could be explained by simple cassette exon inclusion or skipping, potentially at the cost of excluding some inefficiently used MXEs. Among the putative DZF-mediated MXE splicing events represented in two or more MXE datasets, only COL25A1, in which junctions corresponding to only one of the potential MXEs were detected, failed manual inspection. This approach also allowed us to identify regulation of additional genes with MXEs, CASK (Figure 6D) and MAP4K4, both of which have previously been described to have MXEs despite their absence from the MXE catalogs used here (53, 54).

**Figure 6.**
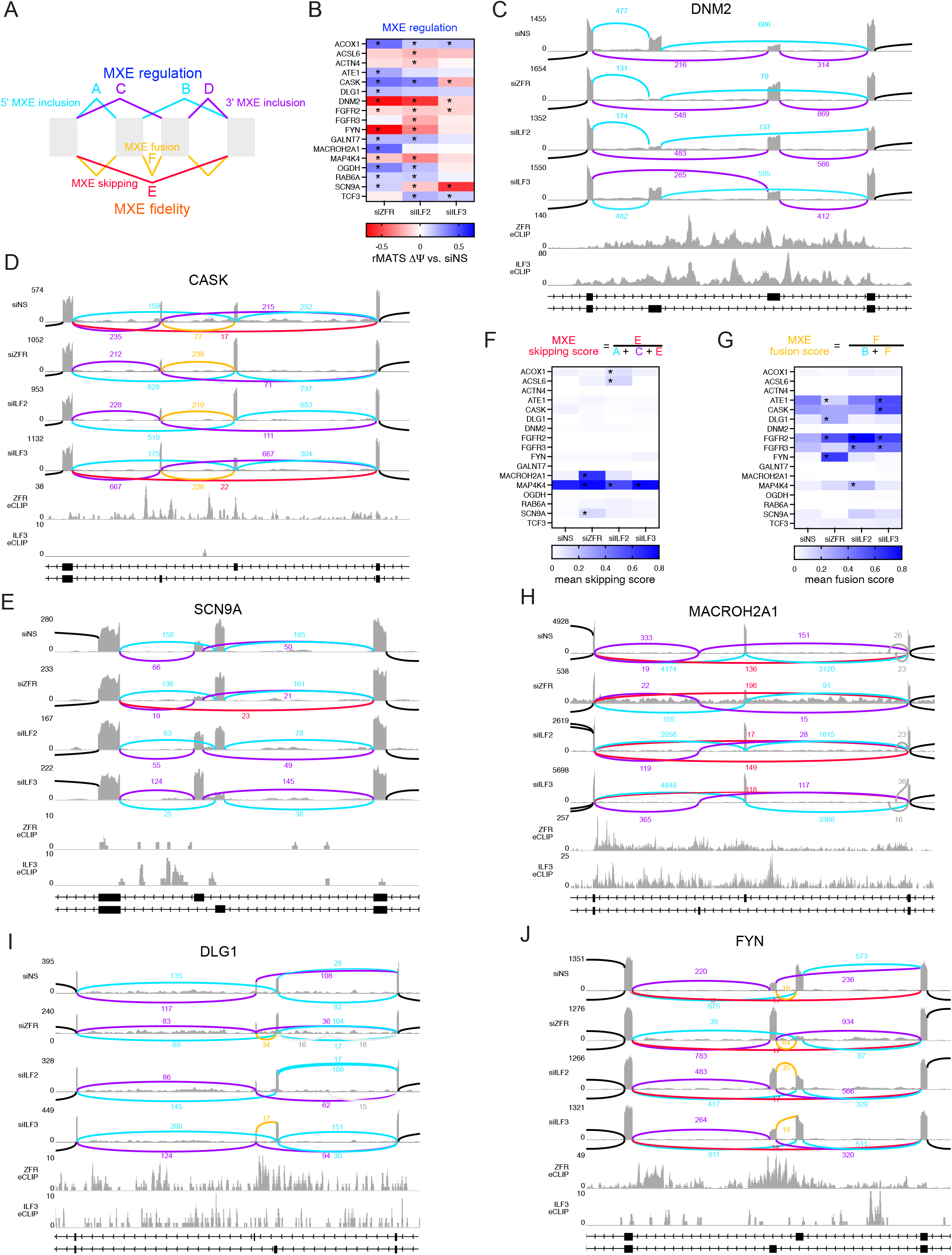
DZF proteins determine regulation and fidelity of MXE splicing. A. Schematic of possible outcomes of alternative splicing of a gene containing two MXEs. Accurate MXE splicing results in pre mRNAs containing junctions A and B or C and D, while failure of MXE splicing fidelity is reflected by skipping (junction E) or fusion (junction F). B. Heatmap of validated MXE splicing events in DZF protein knockdowns relative to siNS control. Asterisks indicate events with rMATS |ΔѰ| > 0.1 and FDR < 0.05. Positive values indicate increased usage of the 5′ exon relative to the 3′ exon in the MXE pair under knockdown conditions. C. Sashimi plot of DNM2 MXE splicing in the indicated knockdown conditions, ZFR eCLIP and Encode ILF3 eCLIP (48) read histograms, and gene model indicating mRNA isoforms resulting from MXE alternative splicing. Mapped reads from three RNA-seq samples per experimental condition were pooled to generate Sashimi plots and junction read counts. Blue lines and read counts correspond to use of the upstream MXE and purple lines and read counts correspond to use of the downstream MXE. Direction of transcription is indicated by chevrons in gene models. Junctions with fewer than 10 reads are not shown. D. Sashimi plot of CASK MXE splicing as in C. Red lines and read counts correspond to isoforms in which all potential MXEs were skipped, and orange lines and read counts correspond to isoforms containing multiple MXEs (fusion events). E. Sashimi plot of SCN9A MXE splicing as in D. F. Heatmap of MXE skipping scores for validated MXE splicing events in DZF protein knockdowns. Statistical significance was determined by two-way ANOVA with Dunnett’s correction for multiple comparisons (n=3; \**P* < 0.05; see Table S4 for fidelity calculations). G. Heatmap of MXE fusion scores for validated MXE splicing events in DZF protein knockdowns. Statistical significance was determined by two-way ANOVA with Dunnett’s correction for multiple comparisons (n=3; \**P* < 0.05; see Table S4 for fidelity calculations). H. Sashimi plot of MACROH2A1 MXE splicing as in D. I. Sashimi plot of DLG1 MXE splicing as in D. J. Sashimi plot of FYN MXE splicing as in D.

As with CEs, ZFR and ILF3 depletion had overlapping but distinct effects on MXE splicing. Overall, the effects of ZFR depletion were more pronounced than those of ILF3 depletion (Figure S8B), with the caveat that these effects could be due to activity of the residual pool of the more abundant ILF3 following knockdown. Providing stronger evidence for differential ZFR and ILF3 functions, we observed cases in which depletion of ZFR and ILF3 had opposite effects on MXE regulation, for example in the CASK and SCN9A genes (Figure 6D and E). Together, these data implicate the DZF proteins as important regulators of mutually exclusive splicing, which can either function together or in opposition to modulate MXE choice.

### Effects of DZF proteins on MXE splicing fidelity

In addition to affecting regulation of MXE splicing by altering exon choice while retaining mutual exclusivity, depletion of the DZF proteins caused loss of MXE splicing fidelity in several genes. Mutually exclusive splicing requires inclusion of one and only one of a set of possible exons into each mRNA. There are two ways in which splicing may deviate from a mutually exclusive pattern, both of which we observed upon depletion of DZF proteins: 1) skipping of all potential exons in the MXE cassette and 2) inclusion of multiple MXEs. We analyzed MXE fidelity in the various DZF protein knockdown conditions by analyzing the relative abundance of junction-spanning reads corresponding to accurate and aberrant MXE splicing (Table S4). For MXE skipping events, we compared the abundance of reads spanning the junction between the constitutive exons upstream and downstream of the MXE cassette to all junction-spanning reads from the upstream constitutive exon 5′ splice site (Figure 6A and F). For MXE fusion events, we normalized reads from the junction between the MXEs to all junction-spanning reads from the upstream MXE 5′ splice site (Figure 6A and G).

We have previously extensively analyzed siZFR-induced skipping of MXEs in MACROH2A1 (Figure 6H), an event which ultimately leads to NMD of the skipped transcript isoform (28). We also observed more moderately enhanced skipping of both MXEs in ACOX1 (ILF2 depletion), ACSL6 (ILF2 depletion), MAP4K4 (ZFR and ILF3 depletion, with the opposite effect in ILF2 depletion), and SCN9A (ZFR depletion; Figure 6E). However, DZF protein depletion also caused inclusion of both exons in a putative MXE pair. We observed elevated levels of MXE fusion transcripts from ATE1 (siZFR and siILF3; Figure 6G), CASK (siILF3; Figure 6D and G), DLG1 (siZFR; Figure 6G and I), FGFR2 (siZFR, siILF2, and siILF3; Figure 6G), FGFR3 (siILF2 and siILF3; Figure 6G), FYN (siZFR; Figure 6G and J), and MAP4K4 (siILF2; Figure 6G). Only one MXE fusion event was suppressed by a DZF protein knockdown, as ATE1 fusion transcripts were less abundant upon siZFR treatment (Figure 6G). Along with double-skipping and double-inclusion, we also observed two cases in which ZFR caused increased usage of a cryptic unannotated exon adjacent to two MXEs: MAP4K4 and DLG1 (Figure 6G and I). These observations suggest that ZFR may have a role in suppressing cryptic splice sites in the process of mutually exclusive exon selection in these genes. Together, these data indicate that the DZF proteins play an important role in not only regulating MXE choice, but also promoting mutual exclusivity.

## Discussion

Here we find that the vertebrate DZF proteins ZFR, ILF2, and ILF3 form a regulatory network that controls many highly conserved and physiologically important splicing events. Using *in vitro* binding assays, we find that the ZFR zinc fingers preferentially bind dsRNA, without apparent sequence preferences. Unexpectedly, eCLIP studies revealed binding of ZFR throughout introns flanking ZFR-regulated CEs and MXEs. Combining our *in vitro* and cellular data, we hypothesize that ZFR initially recognizes one or more dsRNA elements in its pre-mRNA targets and subsequently spreads across introns (Figure 7A). This hypothesis is supported by our finding that many ZFR-regulated exons are flanked by introns that contain multiple conserved sequences with duplex-forming potential (PCCRs; Figure 5A).

**Figure 7.**
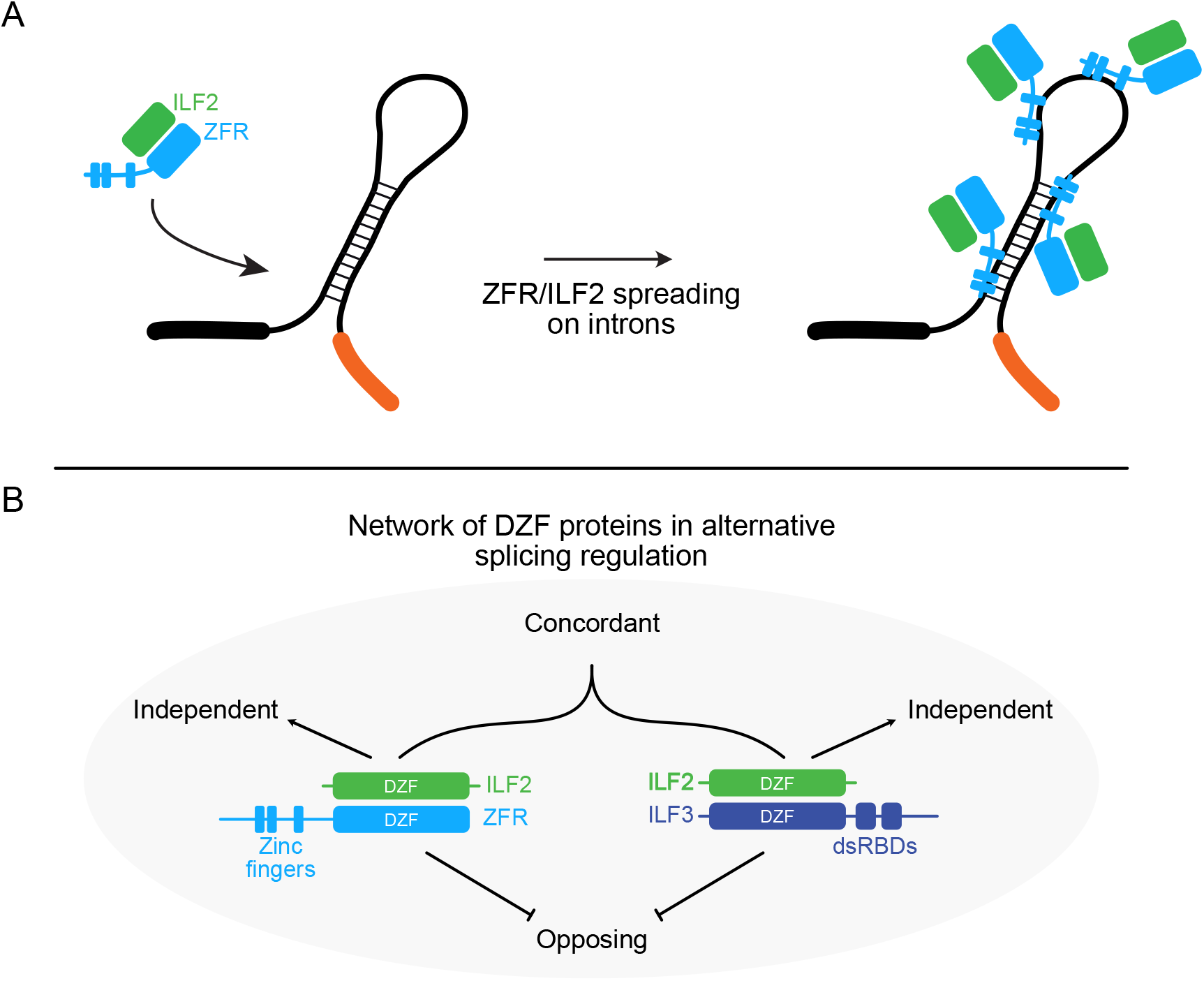
ZFR target recognition and participation in the DZF protein network. A. ZFR associates with broad regions of introns containing dsRNA elements. We propose that ZFR initially recognizes one or more dsRNA segments in its target exons, after which it is able to spread across introns via a currently unknown mechanism. B. The DZF proteins form a physical and functional network in which ILF2 forms mutually exclusive heterodimers with two dsRNA-binding proteins, ZFR and ILF3. ZFR and ILF3 can have independent, concordant, or opposing roles in regulation of CE and MXE splicing.

Unlike most well-studied alternative splicing regulators, both ZFR and ILF3 preferentially interact with dsRNA. ZFR, ILF3, and ADAR proteins have been reported to directly regulate splicing, but the roles and mechanisms of dsRBPs in alternative splicing regulation remain largely unexplored (20, 21, 28, 55). There are several known mechanisms by which RNA structure can modulate splicing efficiency or specificity, all of which may potentially be affected by the DZF and other dsRBPs (56–59). Along with forming interactions with higher-order splicing regulatory complexes (22), ZFR/ILF2 and ILF3/ILF2 heterodimers may modulate dsRNA formation to affect splice site accessibility, trans-acting factor binding to dsRNA or ssRNA, RNA editing, or long-range interactions between splice sites and regulatory elements. Our identification of diverse effects of DZF protein knockdown on CE and MXE alternative splicing, is consistent with highly context-dependent roles for ZFR and ILF3 on dsRNA (Figure 7B).

ILF3 and ZFR both bind dsRNA and form obligate heterodimers with ILF2, but our data indicate that the two proteins have distinct functions in regulation of CE and MXE alternative splicing (Figure 7B). We identify three classes of splicing events that are regulated by DZF proteins: 1) those affected in the same direction and often to a similar extent by depletion of ZFR and ILF3, 2) those altered by depletion of either ZFR or ILF3 but not both, and 3) those regulated in opposing directions by ILF3 and ZFR. Because ILF2 and ILF3 are essential for viability of most or all human cells (60, 61), these experiments required partial depletion via RNAi. This means that we cannot conclusively rule out the involvement of a protein in any given splicing event; however, exons differentially regulated by ZFR and ILF3 tend to have distinct features, suggesting biologically meaningful differences in their biochemical specificity. ZFR positively and negatively regulates exons flanked by long GC-poor introns, while ILF3 promotes inclusion of long exons in high-GC contexts and repression of long exons in low-GC contexts (Figure 4E-H). Together, our results imply that ILF3 and ZFR participate in a complex functional and physical landscape of RNA secondary structure formation and dsRNA-dsRBP interactions.

The targets of DZF-mediated splicing regulation include two particularly interesting and physiologically significant types of exons: ultraconserved CEs and MXEs. Ultraconserved exons are located in several core regulators of RNA splicing, including ZFR (40, 41). While the mechanisms driving such extreme sequence conservation are unclear, ultraconserved exons mediate a dense network of splicing regulation thought to maintain homeostatic control of splicing repressors and activators (42). We now identify ZFR as a participant in the regulatory network of UCE-containing splicing factor genes. Interestingly, our studies suggest that ZFR systematically favors usage of NMD-insensitive isoforms of UCE-containing genes, whether through promotion of exon skipping or inclusion.

Previous studies have identified RBPs that determine regulation or fidelity of individual MXE cassettes (62–64), but to our knowledge the DZF proteins are the first class of RBPs implicated in control of MXE splicing events across multiple genes. Several models for mutually exclusive splicing have been proposed, including incompatibility between introns using the major and minor spliceosomes, post-transcriptional enforcement of MXE splicing by NMD, steric hindrance due to short distances between MXEs, and competition between RNA-RNA duplexes (65). Many MXE splicing events appear to be regulated by the formation of alternative RNA secondary structures, exemplified by the complex Drosophila DSCAM gene, in which a “docking” sequence downstream of constitutive DSCAM exon 5 engages in mutually exclusive interactions with “selector” sequences upstream of exon 6 variants (66, 67).

In light of the evidence that RNA-RNA interactions guide many MXE splicing events, our identification of ZFR, ILF2, and ILF3 as regulators of several important MXE clusters is particularly intriguing. Our data suggest a model in which ZFR and ILF3 use zinc fingers and dsRBDs, respectively, to modulate the formation of RNA duplexes in the course of MXE splicing. In the CASK and SCN9A genes, this results in opposing regulation of MXE splicing upon ZFR and ILF3 depletion (Figure 6D and E). Notably, alternative arrangements of competitive RNA-RNA interactions have been shown to determine MXE regulation and fidelity in ATE1, which we identify here as regulated by ZFR, ILF2, and ILF3 (Figure 6B; 68). ZFR depletion, which caused preferential inclusion of ATE1 exon 7a, phenocopied mutations predicted to prevent a sequence upstream of exon 7a (R1) from base-pairing with a sequence between exons 7a and 7b (R3). The ATE1 R3 sequence was also proposed to engage in alternative interactions with an RNA element immediately upstream of exon 7b (R2). Disruption of R3-R2 base pairing caused elevated fusion of ATE1 MXEs, mirroring the effects of ILF3 depletion. This work thus provides a framework for future investigation of how DZF proteins may interact with and modify RNA secondary structure networks to control alternative splicing regulation and fidelity.

## Acknowledgements

We thank members of the Hogg and Cook groups for helpful suggestions and critical reading of the manuscript and Christopher Burge for advice on RBNS experiments and analysis. High-throughput sequencing was performed in the NHLBI Sequencing and Genomics Core. The Genotype-Tissue Expression (GTEx) Project was supported by the Common Fund of the Office of the Director of the National Institutes of Health, and by NCI, NHGRI, NHLBI, NIDA, NIMH, and NINDS. This work was supported by the Intramural Research Program, NHLBI, NIH, the Wellcome Trust [200898; 203149], and The Darwin Trust of Edinburgh and utilized the computational resources of the NIH HPC Biowulf cluster (http://hpc.nih.gov).

## Author contributions

NH and JRH conceived and designed the study. NH performed eCLIP-seq, RNA Bind-n-Seq, RNA-seq, RT-qPCR, molecular cloning, and IPs; AW expressed and purified recombinant proteins for RNA Bind-n-Seq and EMSA studies and performed EMSA assays. NH, AW, AGC and JRH analyzed data. NH and JRH wrote the manuscript with input from AW and AGC. All authors read and approved the manuscript.

## Supplementary Figure Legends

**Figure S1.**
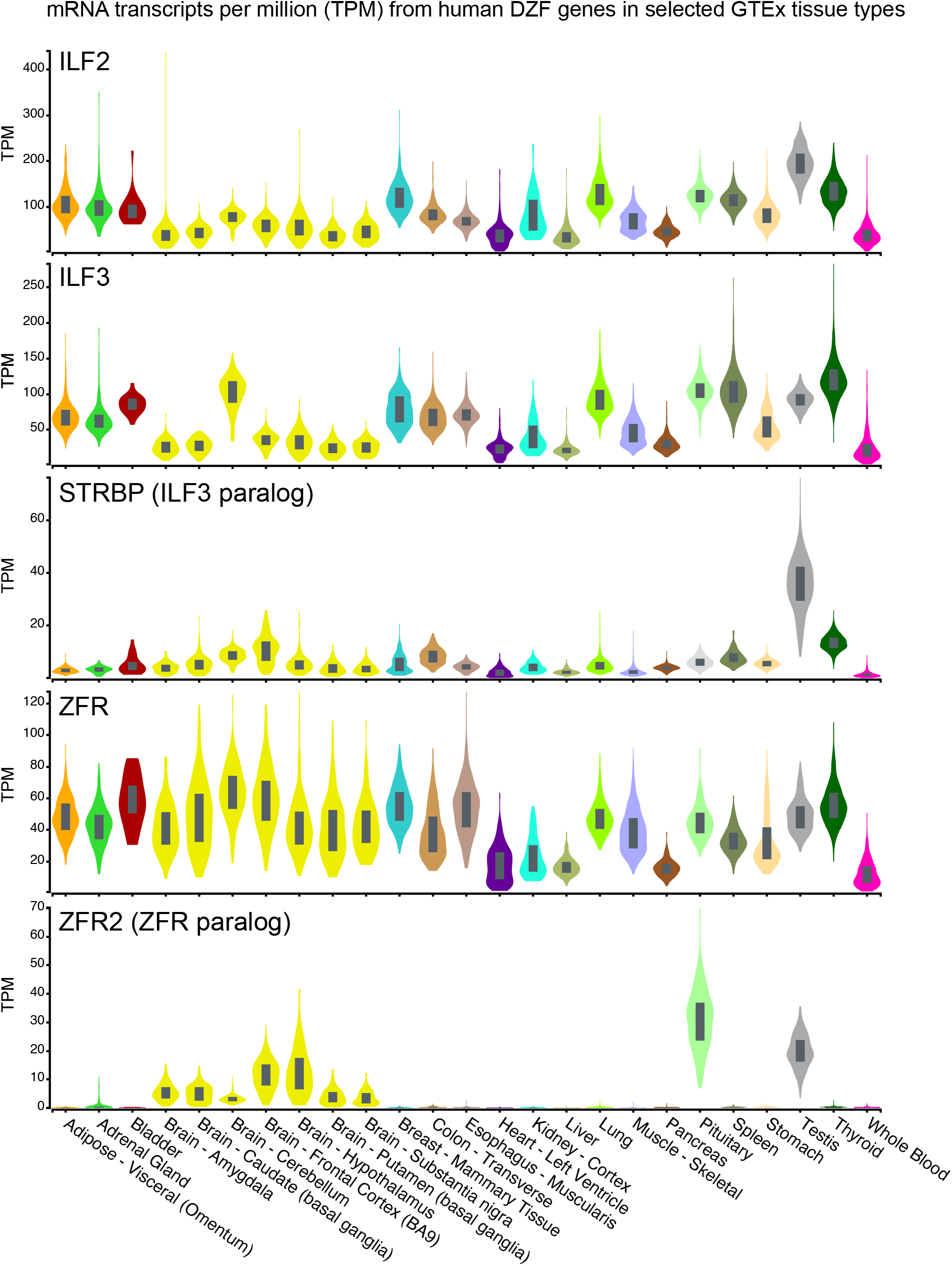
DZF gene expression. A. Relative expression of mRNAs from DZF-containing human genes in the indicated tissue types, as determined and visualized by the GTEx project. The data used for the analyses were obtained from the GTEx Portal on 5/2/2022.

**Figure S2.**
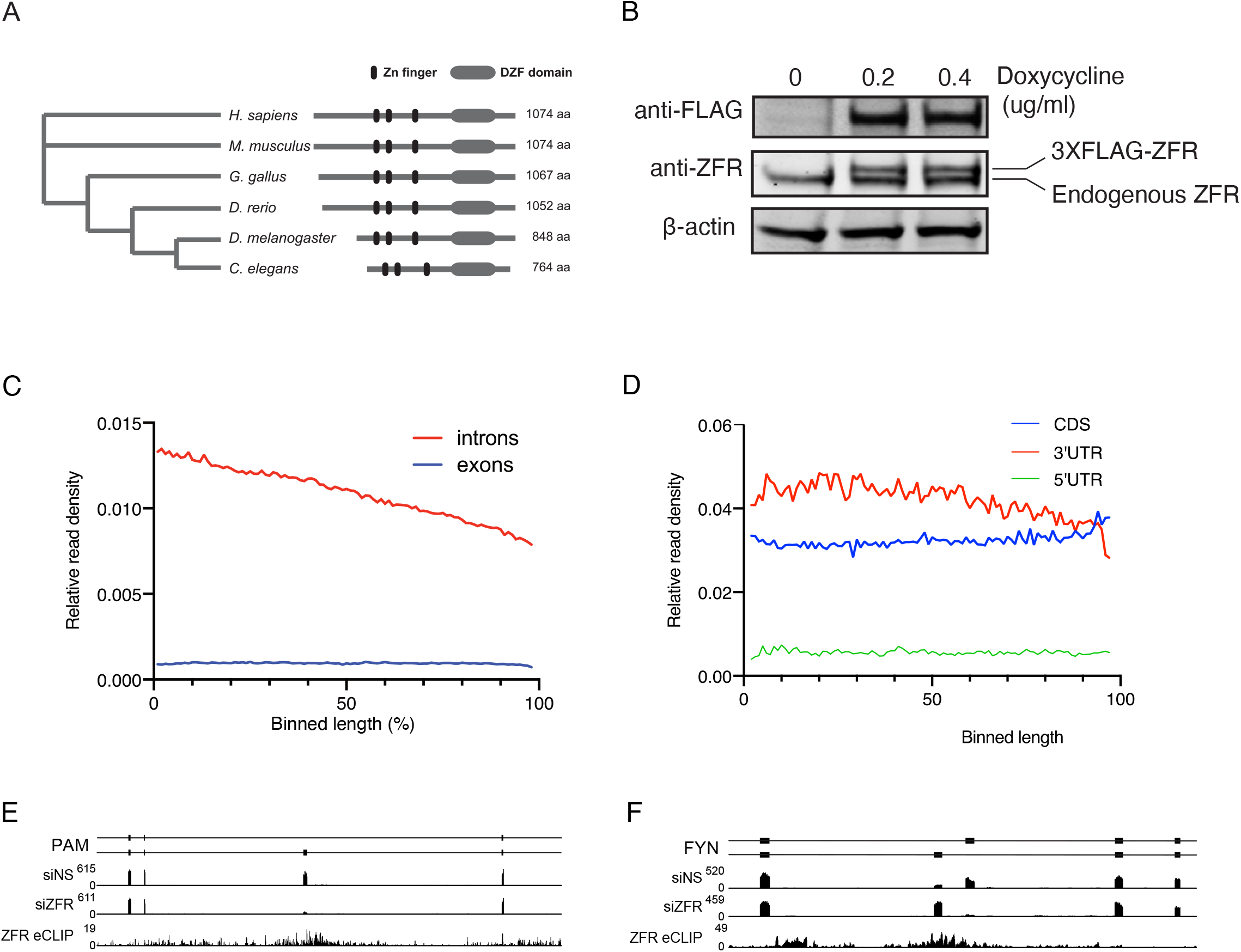
ZFR eCLIP. A. Conservation of ZFR protein architecture among metazoans. Relative positions of zinc-finger and DZF motifs are indicated. B. Immunoblot of HEK-293 Flp-In T-REx cells containing stably integrated 3XFLAG-ZFR expression constructs. Extracts from untreated or doxycycline-treated cells were blotted with the indicated antibodies. C. ZFR eCLIP read densities were computed for exonic and intronic sequences and binned by relative position in annotated introns and exons (72). D. ZFR eCLIP read densities were computed for exonic sequences corresponding to 5′UTRs, 3′UTRs, and coding sequences (CDS) and binned by relative position (72). E. ZFR eCLIP and ZFR and control knockdown RNA-seq read histograms from PAM are shown below gene models corresponding to CE splicing products. Direction of transcription is right to left. F. ZFR eCLIP and ZFR and control knockdown RNA-seq read histograms from FYN are shown below gene models corresponding to MXE splicing products. Direction of transcription is right to left.

**Figure S3.**
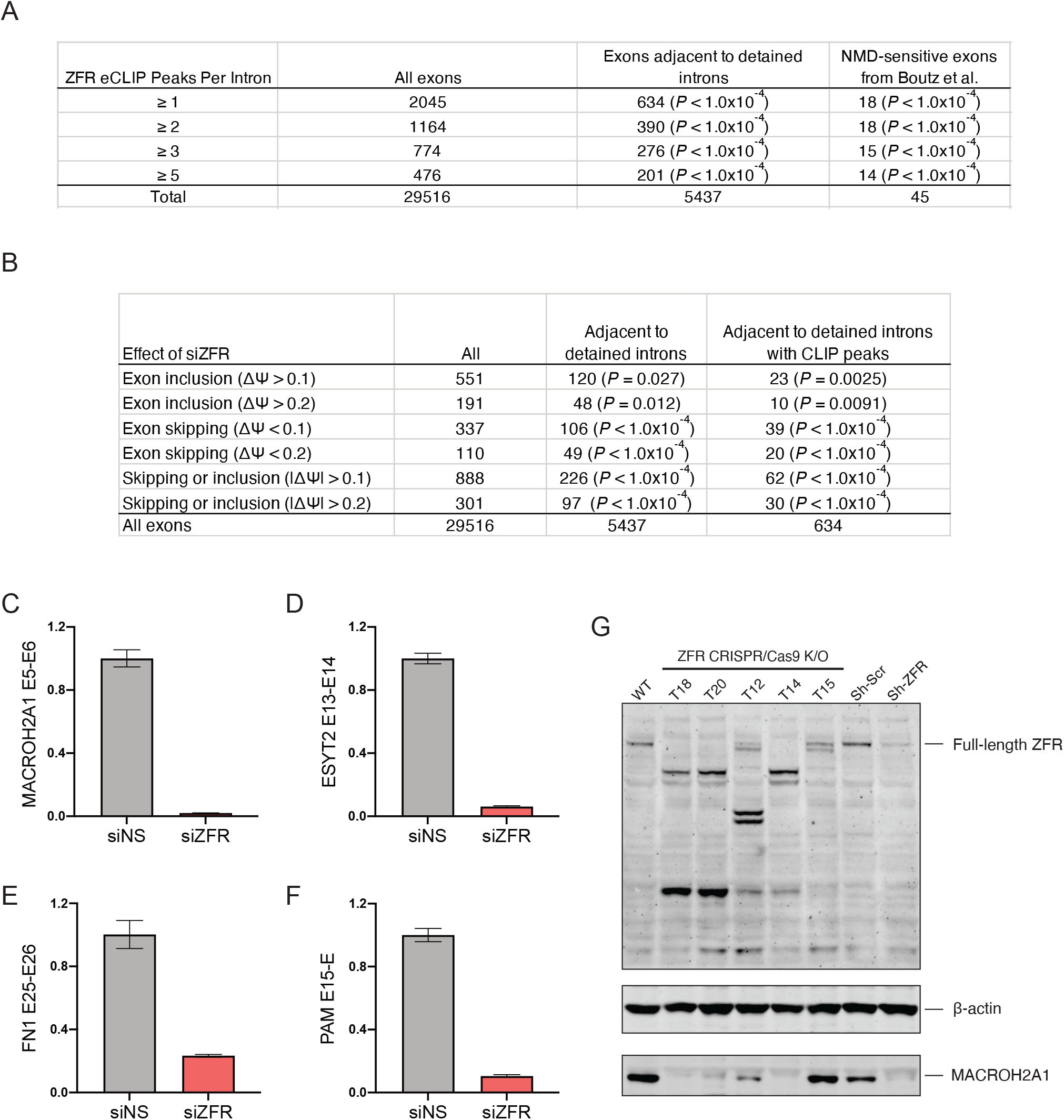
ZFR binding to and regulation of detained introns. A. Table summarizing overlap between intronic ZFR eCLIP peaks and exons associated with detained introns. Empirical *P* values were determined by permutation testing with the RegioneR package (70). B. Table summarizing overlap between rMATS analysis of siZFR vs. siNS knockdown conditions and exons adjacent to detained introns. Empirical *P* values were determined by permutation testing with the RegioneR package (70). C. Isoform specific RT-qPCR of RNA isolated from HEK-293 cells treated with control siRNA or siZFR to assess alternative splicing of MACROH2A1 mRNAs. Relative levels of mRNA isoforms containing the indicated exon junction were determined by normalization to GAPDH mRNA (n=3, error bars indicate SD). D. Quantification of ESYT2 alternative splicing as in C. E. Quantification of FN1 alternative splicing as in C. F. Quantification of PAM alternative splicing as in C. G. Immunoblotting of extracts from parental and clonal CRISPR/Cas9 ZFR knockout HEK-293 cells, with extracts from cells treated with control or anti-ZFR shRNAs as controls. ZFR expression was detected with an antibody raised against the C-terminus of the protein (28). Disruption of the splicing regulatory function of ZFR was verified by immunoblotting for MACROH2A1, which depends on ZFR for expression. Clone T18 was used for further experiments.

**Figure S4.**
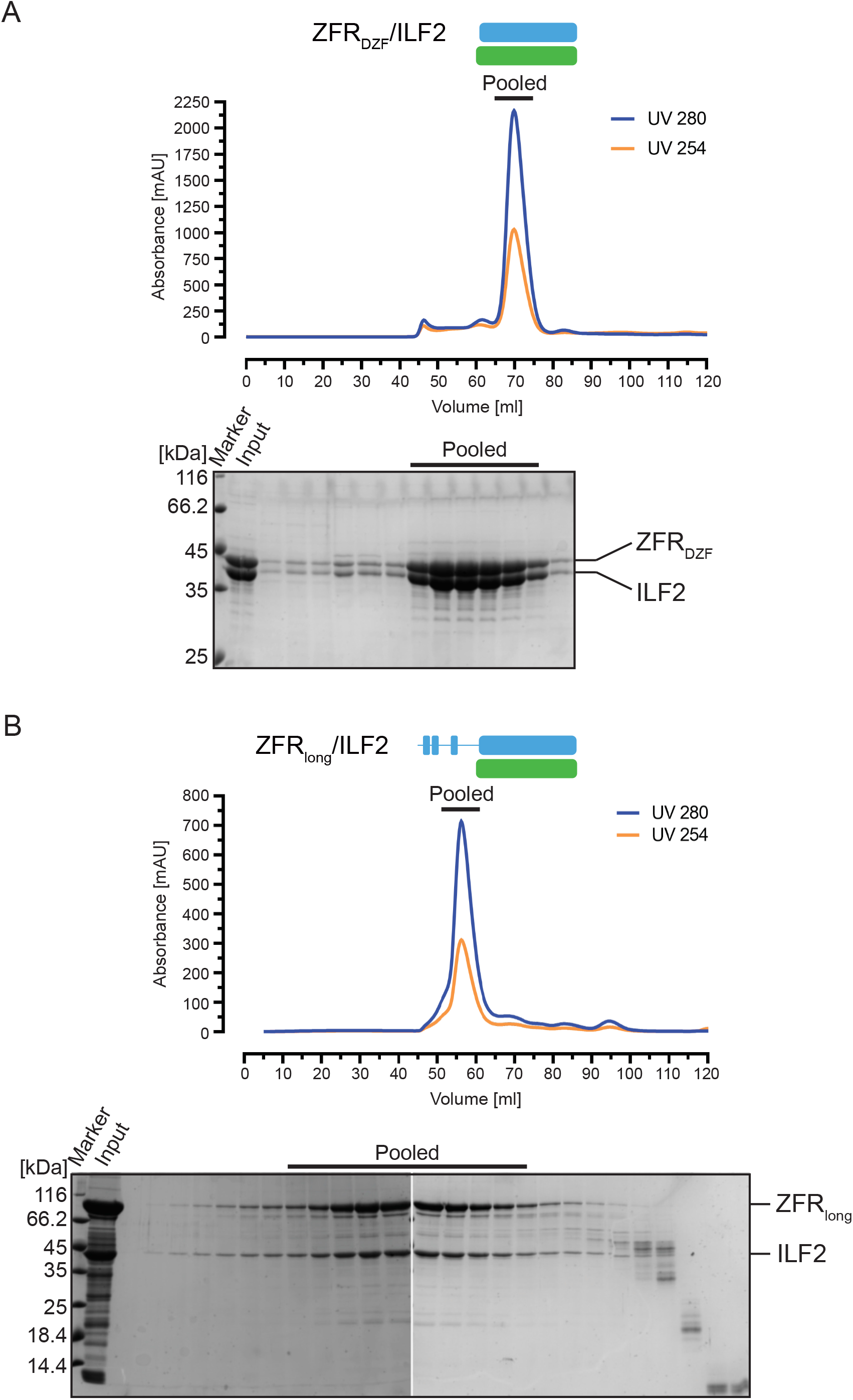
ZFR-ILF2 complex purification. A. Characterization of co-expressed and purified ZFR_DZF_/ILF2 complexes. Top, chromatogram of ZFR_DZF_/ILF2 elution from a Superdex S200 size exclusion column. Bottom, SDS-PAGE of fractions eluted from the S200 column. The indicated fractions were pooled for further experiments. B. As in A, with co-expressed ZFR_long_/ILF2.

**Figure S5.**
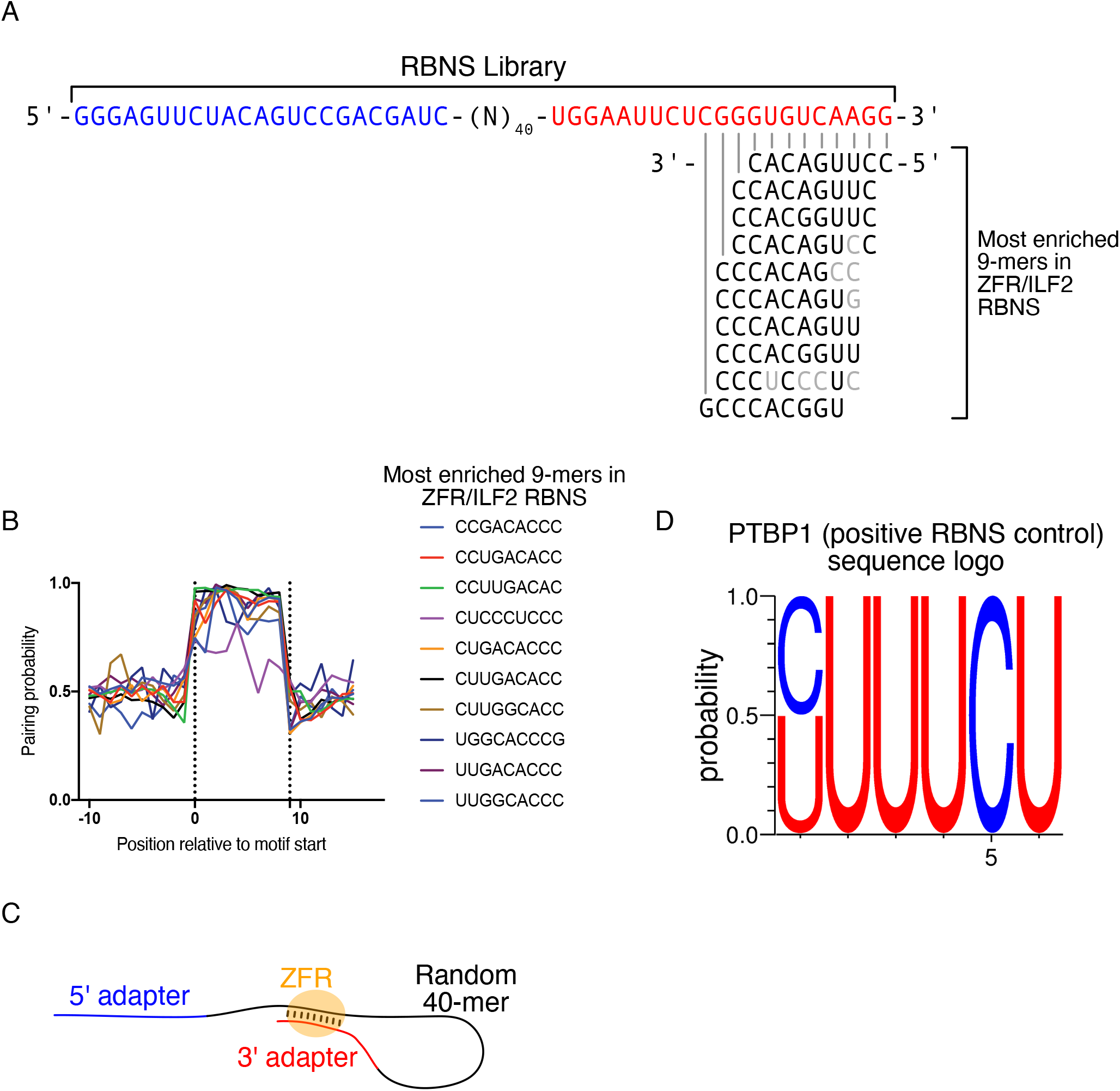
ZFR RNA Bind-n-Seq recovers sequences complementary to PCR primer binding sites. A. Schematic of RBNS library. 5′ and 3′ adapters for PCR primer binding are indicated in blue and red, respectively, flanking a 40 nt random sequence. The most highly enriched 9-mers in ZFR_long_/ILF2 RBNS are fully or partially complementary to a sequence at the end of the 3′ primer binding site. B. Predicted pairing probabilities for most highly enriched 9-mers in ZFR RBNS. Potential for pairing to sequences found in all RNAs in the pool leads to high base-pairing probability for ZFR-enriched sequences. C. Schematic of predicted pairing between 3’ primer binding site and RBNS library sequences highly enriched with ZFR. D. Sequence logo of most highly enriched 6-mers recovered in positive control PTBP1 RBNS.

**Figure S6.**
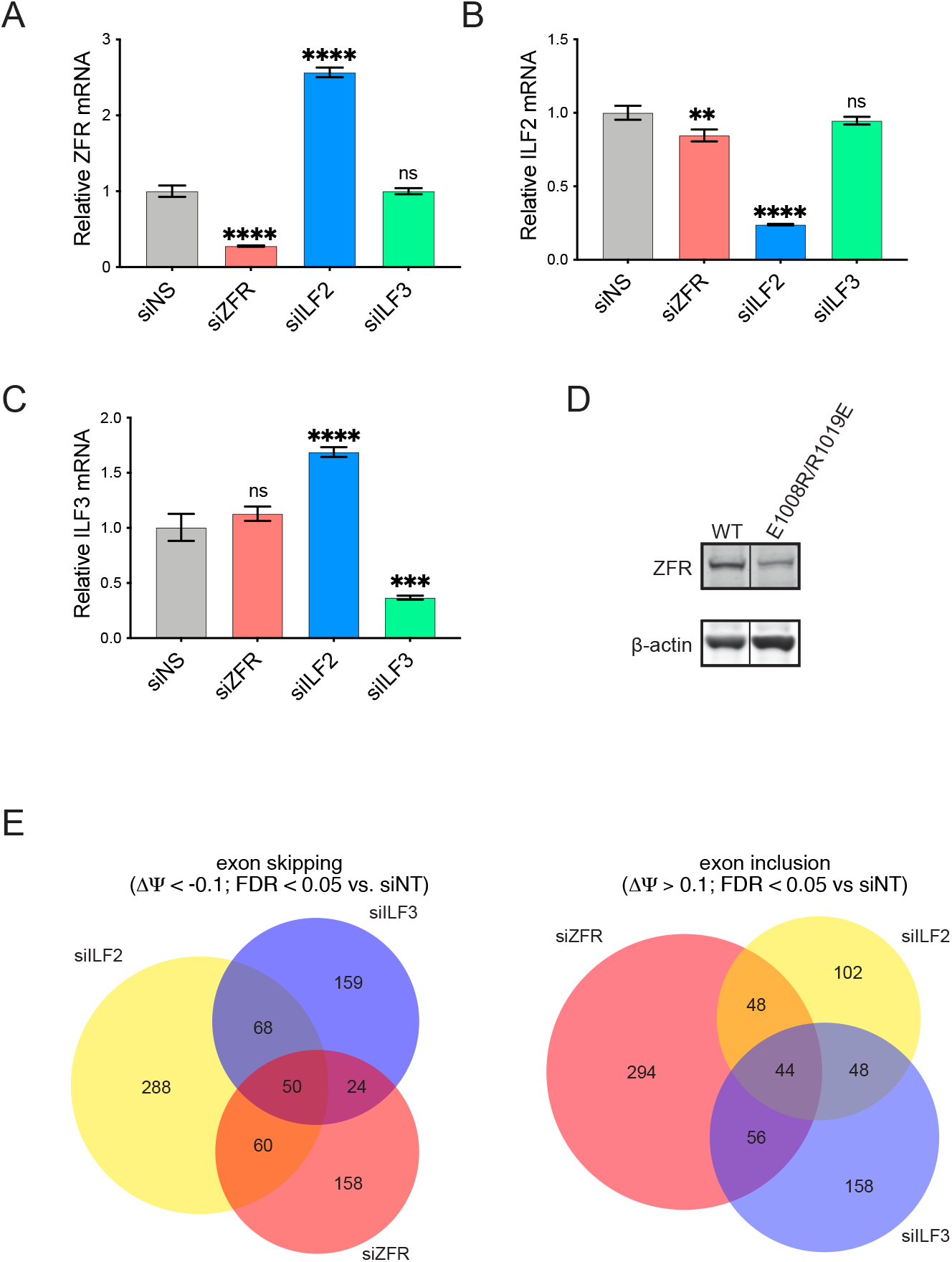
Effects of DZF protein depletion on DZF mRNA abundance and alternative splicing. A. RT-qPCR of total endogenous ZFR mRNA in the indicated knockdown conditions. Statistical significance was determined by two-way ANOVA, with Sidak’s correction for multiple comparisons (n = 3; error bars indicate ±SD; * *P* < 0.05; ** *P* < 0.01; *** *P* < 0.001; **** *P* < 0.0001). B. RT-qPCR of total endogenous ILF2 mRNA as in A. C. RT-qPCR of total endogenous ILF3 mRNA as in A. D. Immunoblot of 3XFLAG-ZFR proteins expressed from transgenes stably integrated in HEK-293 Flp-In T-REx cells upon treatment with doxycycline (1 µg/ml). The E10008R/R1019E mutation is predicted to abolish ZFR binding to ILF2. E. Venn diagrams of CE skipping (left) and inclusion (right) identified by rMATS analysis of the indicated DZF protein knockdowns.

**Figure S7.**
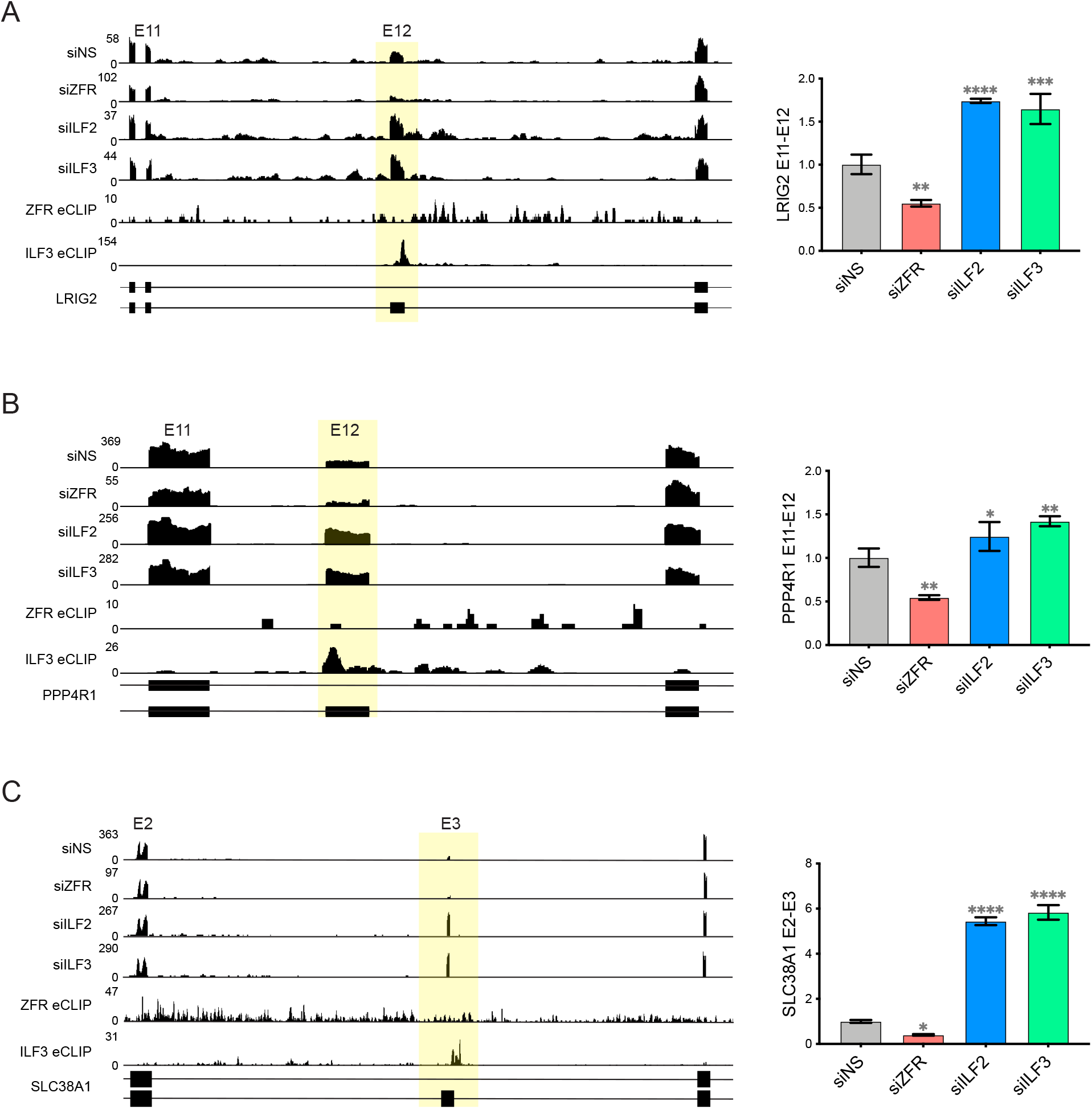
Opposing effects of ZFR and ILF3 depletion on specific CE alternative splicing events. A-C Left, ZFR eCLIP, Encode ILF3 eCLIP (48) and DZF knockdown RNA-seq read histograms from the indicated genes, with gene models indicating mRNA isoforms produced by alternative CE splicing. Direction of transcription is left to right. Right, RT-qPCR analysis of the relative abundance of mRNAs containing the indicated splice junctions, normalized to GAPDH mRNA levels. Statistical significance was determined by two-way ANOVA, with Sidak’s correction for multiple comparisons (n = 3; error bars indicate ±SD; * *P* < 0.05; ** *P* < 0.01; *** *P* < 0.001; **** *P* < 0.0001).

**Figure S8.**
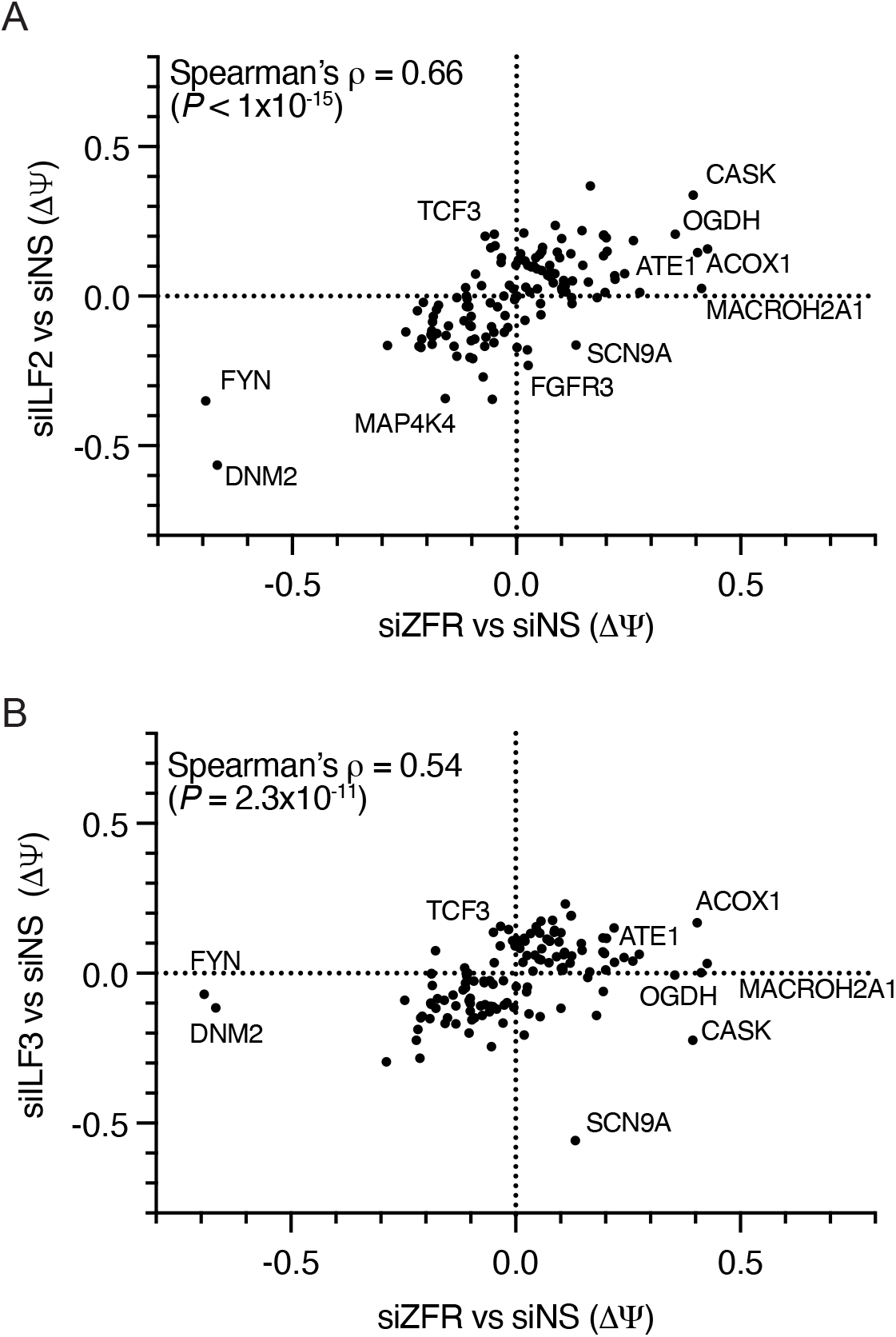
Effects of DZF protein depletion on MXE alternative splicing. A. Scatterplot comparing response of MXE events meeting rMATS read count cutoffs and regulated in at least one DZF protein knockdown condition (|ΔѰ| > 0.1 and FDR < 0.05) to depletion of ZFR or ILF2. Positive values indicate increased usage of the 5′ exon relative to the 3′ exon in the MXE pair under knockdown conditions. B. As in A, comparing the effects of ZFR and ILF3 depletion.

## Supplementary Tables

Table S1. Compiled rMATS data (ZFR, ILF2, and ILF3; all event types)

Table S2. RBNS results (6-mers and associated B values)

Table S3: MXE catalog (compiled genes; compiled genes with rMATS coverage)

Table S4: MXE fidelity calculations

Table S5: Oligonucleotide sequences

## Materials and Methods

### Cell culture

HEK-293 Tet-Off (HEK-293TO) cells (Clontech) and HEK-293 Flp-In T-REx cells (ThermoFisher Scientific) were maintained in Dulbecco’s Modified Eagle’s Medium (DMEM; ThermoFisher Scientific) supplemented with 10% FBS (Gibco), 100 U/mL of penicillin, 100 µg/mL streptomycin (Gibco), and 0.3 mg/mL of L-glutamine (Gibco), at 37°C and 5% CO_2_. Cell lines were periodically tested for mycoplasma contamination using the MycoAlert Mycoplasma Detection Kit (Lonza).

### Plasmids

Full-length cDNAs of ILF2 and ILF3 were PCR amplified from cDNA clones (Dharmacon) and inserted into pcDNA5/FRT/TO (ThermoFisher Scientific) downstream of a 3XFLAG tag previously inserted into the vector (28) to make the resulting plasmids pcDNA5/FRT/TO-3XFL-ILF2 and pcDNA5/FRT/TO-3XFL-ILF3.

### Alphafold-Multimer

Alphafold version 2.1.1 was used to predict the structure of the heterodimer between the DZF domains of ZFR (Uniprot ID Q96KR1) and ILF2 (Q12905), using the multimer model preset (69).

### RNAi

siRNA mediated depletion of gene expression was performed using Silencer Select siRNAs (ThermoFisher Scientific) as previously described (28). Briefly, 1.5X10^5^ cells were reverse transfected with 120 pmol of siRNAs in 6-well plates using Lipofectamine RNAiMAX (ThermoFisher Scientific). Cells were harvested for RNA extraction 72 hours after transfection. The following siRNAs were used: si-ZFR: 5′-GCACUUAAAAGGGCGAAGATT-3′; siILF2: 5′-CAGGGACAUUUGAAGUGCATT-3′; siILF3: 5′-AGAAGACAAGUACGAAAUATT-3′; si-NS: 5′-UAAGGCUAUGAAGAGAUAC-3′).

### Mammalian cell line generation

Human Flp-In T-REx-293 cells expressing 3×FLAG-ILF2, ILF3, and ZFR were generated following the manufacturer’s protocol (ThermoFisher Scientific). Flp-In T-REx-293 cells were transfected with pcDNA5/FRT/TO-3XFL-ILF2, pcDNA5/FRT/TO-3XFL-ILF3, or pcDNA5/FRT/TO-3XFL-ZFR and pOG44 plasmids (ThermoFisher Scientific) with TurboFect transfection reagent according to the manufacturer’s protocol (ThermoFisher Scientific). Cells were selected and maintained with hygromycin (100 μg/mL; Invitrogen) 10–14 days after transfection. Polyclonal cells were treated with doxycyline hyclate (1 μg/mL; ThermoFisher Scientific) for 48–72 hr for induction of transgene expression. Transgene expression was confirmed by western-blot analysis. For CRISPR/Cas9-mediated gene knockout of ZFR in HEK-293TO cells, plasmids containing pre-designed guide RNAs (sc-407242) and homology-directed DNA repair templates harboring puromycin resistance and RFP coding sequences (sc-407242-HDR) were purchased from Santa Cruz Biotechnology. 1.5X10^5^ cells were transfected with 1 μg of each plasmid using TurboFect transfection reagent (Thermo Scientific). Medium was replaced after 24 hr, and cells were selected with puromycin (2.5 μg/mL) after 72 hr. Single cells were sorted into 96-well plates based on RFP expression for clonal expansion.

### RNA isolation and RT-qPCR

RNA was extracted using the RNeasy Mini Kit (Qiagen), with on-column DNase digestion with the RNase-free DNase Set (Qiagen). RNA concentration and purity were evaluated using a NanoDrop One spectrophotometer (ThermoFisher Scientific). cDNA was synthesized from 500 ng of total RNA using the Maxima First Strand cDNA synthesis kit (ThermoFisher Scientific). cDNA was diluted to 1:20 with water, and 3 µl was used for qPCR with iTaq Universal SYBR Green Supermix (BioRad). Relative levels of RNA were calculated by the ΔΔCt method using GAPDH as an internal control. At least three biological replicates were performed, from which average and standard deviation values were calculated. Oligonucleotide sequences are provided in Table S5.

### Cell extract preparation and immunoprecipitation (IP)

Cell extract preparation and immunoprecipitation were performed based on previously described methods (73). Flp-in T-TEx cells expressing FLAG-tagged ILF2, ILF3 or GFP were seeded in 15 cm plates and induced with doxycycline hyclate (200 ng/mL) for 72 hr. After trypsinization, cells were washed with cold PBS and transferred to 1.5 mL tubes. Cells were centrifuged at 300 g for 5 min, and cell pellets were resuspended in 5X packed cell volume of cold lysis buffer (20 mM HEPES, pH 7.5, 2 mM MgCl_2_, 10% glycerol, 1 mM DTT, 1 mM EGTA, 0.1% NP40, Halt protease and phosphatase inhibitor (EDTA free; ThermoFisher Scientific) and kept on ice for 5 min. Cells were homogenized in a Dounce homogenizer using a tight pestle, and NaCl was added dropwise to a final concentration of 150 mM. Cell lysis was checked under a light microscope. Following confirmation of >95% cell lysis, homogenates were transferred to pre-chilled 1.5 mL tubes and rocked for 30 min in 4° C. Cell lysates were cleared by centrifugation at 12,000 g for 10 min at 4° C, and supernatants were transferred to new tubes, snap frozen in liquid nitrogen, and stored at −80°C.

For immunoprecipitation (IP), lysates were thawed on ice and centrifuged at 12,000 rpm for 5 min at 4°C to remove any precipitated material, and equal amounts of protein were used for each IP. Pierce IgG magnetic beads (25 µL; ThermoFisher Scientific) conjugated with anti-FLAG-M2 monoclonal antibody (10 µg; Sigma-Aldrich) were mixed with lysates and rotated in a cold room for 1.5 hr. Beads were washed with 500 µL of lysis buffer supplemented with 150 mM NaCl without glycerol and resuspended in 500 µL of MNase buffer (50 mM Tris HCl, pH 7.5, 2 mM CaCl_2_, 100 mM NaCl, 2.5 mM MgCl_2_). Samples were split in half for MNase (2000 units, NEB) treatment at 27°C for 30 min or mock treatment on ice. After 30 minutes, MNase was inactivated by adding 12 µL of 0.5 M EGTA, and all samples were washed twice with 500 µL of lysis buffer supplemented with 150 mM NaCl without glycerol. After the last wash, tubes were briefly spun to allow removal of residual buffer, and proteins were eluted with 50 µL of 1x NuPAGE LDS buffer (ThermoFisher Scientific) by heating at 70°C for 10 min. Purified proteins were transferred to new tubes, supplemented with 100 mM DTT, and stored at −30°C before separation on 4-12% NuPAGE Bis-Tris protein gels (ThermoFisher Scientific).

### Protein expression and purification

Two constructs of recombinant human ZFR in complex with ILF2_29-390_ were prepared: ZFR_DZF_ (residues 725-1074, dimer is denoted as ZFR_DZF_/ILF2) and ZFR_long_ (residues 317-1074, dimer is denoted as ZFR_long_/ILF2). ZFR and ILF2 constructs were co-expressed as N-terminal hexahistidine-tagged proteins in *E. coli* B834(*DE3*). Cells were grown in 2xTY at 37 °C and cooled to 18°C when an OD_600_ of 0.8 was reached. Protein expression was induced with 0.2 mM IPTG, at 18°C overnight. Cells were lysed using a cell disruptor (Constant Systems) in 20 mM Tris-HCl pH 8.0, 150 mM NaCl, 10 mM imidazole pH 8.0, 0.5 mM β-Mercaptoethanol (β-ME), supplemented with a protease inhibitor cocktail (Roche) and DNAse I (Sigma Aldrich). Lysates were cleared by centrifugation (50,000g, 4 °C, 45 min) and bound in batch to Ni-NTA resin (Cytiva), pre-equilibrated in lysis buffer, at 4°C for 2 h. Beads were packed into a column, washed with lysis buffer and eluted over a linear gradient of 10 to 500 mM imidazole in buffer (20 mM Tris-HCl pH 8.0, 150 mM NaCl, 0.5 mM β-ME). Protein complexes were dialysed into 20 mM HEPES pH 7.5, 50 mM NaCl, 1 mM DTT and further purified using heparin sepharose chromatography. Samples were eluted using a linear gradient of 50 to 1000 mM NaCl. For ZFR_long_/ILF2 an additional cation exchange step was included. ZFR_long_/ILF2 was dialysed into 20 mM HEPES pH 7.5, 50 mM NaCl and 1 mM DTT overnight, applied to a 6 ml S column, washed, and eluted with a 50 to 1000 mM NaCl gradient. Finally, both samples were concentrated and applied to a HiLoad® 16/600 Superdex® 200 pg equilibrated in size-exclusion buffer (20 mM HEPES pH 7.5, 50 mM NaCl and 1 mM DTT). Peak fractions were pooled, concentrated and flash-frozen and stored at −80 °C until use.

### Electrophoretic mobility shift assays

RNA oligonucleotides were purchased from Biomers.net (Ulm, Germany) with either a DY681-or Cy5-fluorophore attached to the 5′ end. Oligonucleotides were reconstituted in water. Complementary strands were mixed and heated to 95°C with cooling overnight to RT to form duplexes. Binding reactions contained protein and RNA or RNA:DNA oligos in 20 mM HEPES pH 7.5, 150 mM potassium acetate, 4 mM magnesium acetate and 1 mM DTT. EMSAs used either ZFR_DZF_/ILF2 complex or ZFR_long_/ILF2 with identical conditions. Each reaction was assembled in 10 µl, containing 0.5 µM RNA or RNA:DNA and increasing concentrations of protein. Samples were incubated on ice for 1h. A 20 cm × 20 cm 8 % native polyacrylamide gel in 0.5× TBE buffer was pre-run at 2W at 4°C for 1 h. To each sample, 2 µl 6× native gel loading buffer (50 % (v/v) glycerol and 0.25 % (w/v) bromophenol blue) was added. 4 µl of each reaction was loaded onto the gel and run at 2 W at 4 °C for 1 h. Migration of fluorescently labeled RNA oligos was visualized using the Odyssey® CLx (LI-COR) imaging system at 700 nm. Images were converted to grayscale using Image Studio.

### Immunoblot analysis

RIPA buffer (10 mM Tris-HCl pH 8.0, 1 mM EDTA, 0.5 mM EGTA, 150 mM NaCl, 1% NP40, 0.1% sodium deoxycholate, 0.1% SDS, and Halt Protease Inhibitor cocktail (Thermo Scientific)) was used for cell lysis. Following centrifugation at 9000 g, cell lysate protein concentrations were measured using Pierce 660 nm Protein Assay Reagent (ThermoFisher Scientific) on a multimode plate reader (Infinite M200 PRO, Tecan). Equal amounts of protein were separated on 4-12% Nupage Novex Bis-Tris protein gels (ThermoFisher Scientific). For ZFR immunoblots, gels were soaked in 2X NuPAGE transfer buffer (ThermoFisher Scientific) supplemented with 0.03% SDS for 20 min before transfer. Proteins were transferred to 0.45 μM Nitrocellulose membranes (Bio-Rad) with an XCell II Blot Module (ThermoFisher Scientific). Membranes were blocked overnight with Blocking Buffer for Fluorescent Western Blotting (Rockland), and blotted with primary antibodies against ZFR (Bethyl Laboratories, at 1:1000), ILF2 (Bethyl Laboratories, at 1:5000), ILF3 (Bethyl Laboratories, at 1:500), β-actin (Cell Signaling, 1:1000). Fluorescently labeled secondary antibodies (ThermoFisher Scientific) were used at 1:10,000. Membranes were scanned with either an Odyssey Imaging System (LI-COR) or Amersham Typhoon 5 Gel and Blot Imaging System (GE Healthcare).

### eCLIP-seq

eCLIP was performed based on the protocol in (35). As in irCLIP (36), adapters (Integrated DNA Technologies) were labeled with IRdye-800CW-DBCO (Li-Cor) as described to allow visualization of adapter-ligated RNAs (74). 3XFLAG-ZFR and 3XFLAG-PTBP1 expression was induced for 72 hours by addition of 0.4 µg/mL doxycycline hyclate (ThermoFisher Scientific) prior to cell lysis. ZFR eCLIP library preparation, sequencing, and data processing were performed at the same time and identically to PTBP1 eCLIP reported previously (74). Briefly, fastq files were trimmed with Cutadapt (75), mapped against human RepBase sequences to remove sequences mapping to repetitive elements with STAR aligner (76, 77). The remaining reads were mapped to human genome hg19 with STAR aligner. ZFR and PTBP1 eCLIP peaks were called with Clipper software with parameters --bonferroni -- superlocal --threshold-method binomial and normalized to size-matched input controls as described (35, 48, 78). Peaks with Clipper *P* < 10^-11^ were used for analysis of intersections with significant rMATS splicing events, detained introns (38), and PCCRs (46) using the bedtools intersect function (80). CLIPSeqTools options distribution_on_genic_elements and distribution_on_introns_exons were used to compute eCLIP read density across gene features, using annotation files provided by the authors (72). Metagene analyses were performed with the rMAPS server, using rMATS SE JCEC output files and ZFR eCLIP Clipper peak output files (71). eCLIP coverage maps were generated using Integrative Genomics Viewer (79).

Using a search space of introns from highly expressed genes, LOLA software (47) was used to assess the overlap between PCCRs in which complementary regions were separated by no more than 1000 nt (46) and ZFR and PTBP1 eCLIP peaks (this study; 74) or eCLIP peaks from the complete catalog of ENCODE studies in HepG2 and K562 cells (48). To minimize confounding effects of transcript abundance, genes with RPKM values in the top quartile of in siNS control RNA-seq were selected for further analysis of eCLIP peak overlap. Introns were extracted from Encode GRCh37 gene annotations and redundant introns were removed using bedtools merge function (80). Introns were divided into 200 bp fragments for LOLA analysis of overlap between eCLIP peaks and PCCRs.

### RNA Bind-n-Seq

RBNS of 6X-His tagged ZFR_DZF_/ILF2, ZFR_long_/ILF2, and PTBP1 was performed based on established protocols (43, 44), with modifications. The DNA template for T7 transcription of the RBNS library containing 40 random nucleotides was synthesized by Integrated DNA Technologies and transcribed with the MEGAshortscript T7 transcription kit (ThermoFisher Scientific). Four concentrations of each ZFR/ILF2 complex (5, 20, 80, and 320 nM) or two concentrations of control protein PTBP1 (20 and 80 nM) were equilibrated in 45 µl binding buffer (20 mM HEPES pH 7.5, 150 mM potassium acetate, 3 mM magnesium acetate, 0.01% Tween-20, 1 mM DTT, 1 mM zinc sulfate, 0.5 mg/ml BSA) for 30 minutes at RT, and 90 µl 1 µM RBNS RNA library supplemented with 20 units of SUPERaseIn (ThermoFisher Scientific) was added and rotated for 1 hr at RT. Pre-washed 25 µl His-Tag Dynabeads (ThermoFisher Scientific) were added, rotated for 1 hour at RT, and washed three times with wash buffer (20 mM HEPES pH 7.5, 150 mM potassium acetate, 0.01% Tween, 0.5 mM EDTA, 0.5 mg/ml BSA). Bound RNA was eluted with 100 µl elution buffer (10 mM Tris, pH 7.0, 1 mM EDTA, 1% SDS) by heating at 70°C for 10 min. Eluted RNA was purified with Zymo RNA Clean & Concentrator-5 columns (Zymo Research), reverse-transcribed with SuperScript IV (ThermoFisher Scientific), PCR amplified with AccuPrime PFX SuperMix (ThermoFisher Scientific), and sequenced on the Illumina HiSeq 3000 platform. Libraries from 0.5 pmol of the starting RBNS RNA pool and RBNS control reactions lacking recombinant RBPs were generated as input and negative control samples, respectively. RBNS sequencing data were analyzed using RBNS pipeline software (https://github.com/alexrson/rbns_pipeline; 44) and the Vienna RNAfold package (81). Sequences complementary to sample barcodes were excluded from analyses. For pairing probability analyses, the 9-mers with the largest RBNS B-factors in the 320 nM ZFR_long_/ILF2 condition were assessed.

### RNA-seq and alternative splicing analysis

RNA-seq was performed as previously described (28). Total RNA (1 µg) was used for cDNA library preparation. Ribo-Zero rRNA Removal Kit (Epicenter) and Illumina TruSeq Stranded Total RNA Sample Preparation Kit (Illumina) were used for depletion of ribosomal RNA, and cDNA library synthesis, respectively. Library quality was assessed on an Illumina MiSeq system and sequenced on the Illumina HiSeq 3000 platform to generate 75 bp paired-end reads. Sequencing quality was assessed with fastqc (82), and raw reads were aligned to the human genome assembly hg19 using Hisat2 and the combined genome and transcriptome index provided by the authors, with novel junction alignments allowed (83).

Alternative splicing analysis was performed with rMATS software and Ensembl GRCh37 gene annotations (84). rMATS output files were stringently read-count filtered to include only events in which junctions corresponding to either the included or excluded product were represented by an average of at least 33 reads per sample in one or more experimental conditions, and redundant CEs were removed using the bedtools merge function (80). Unless otherwise noted, |ΔѰ| > 0.1 and FDR < 0.05 were used to identify significant alterations in splicing. Sashimi plots were generated using Integrative Genomics Viewer (79) and annotated using Affinity Designer software (Serif), and proportional Venn diagrams were constructed using DeepVenn (85).

Analysis of overlap between DZF protein-regulated alternative splicing and sequence features or eCLIP peaks was performed using rMATS event coordinates following read-count filtering. Redundant exonic or intronic intervals were removed using the bedtools merge function (80). The RegioneR package was used for permutation analysis to derive empirical *P* values (10,000 iterations for all analyses shown), with resampleRegions and numOverlaps options (70). Coordinates of detained introns were obtained from (38) and were filtered to remove introns containing snoRNAs in Gencode v39 annotations (86). Sequence compositions of introns and exons represented in rMATS were determined with the bedtools nuc tool (80).

